# Microglia and perivascular macrophages act as antigen presenting cells to promote CD8 T cell infiltration of the brain

**DOI:** 10.1101/2021.04.09.439241

**Authors:** Emma N. Goddery, Cori E Fain, Chloe G Lipovsky, Katayoun Ayasoufi, Lila T. Yokanovich, Courtney S. Malo, Roman H. Khadka, Zachariah P. Tritz, Fang Jin, Michael J. Hansen, Aaron J. Johnson

**Affiliations:** Department of Immunology, Mayo Clinic, Rochester, MN, USA; Mayo Clinic Graduate School of Biomedical Sciences, Mayo Clinic, Rochester, MN, USA; Department of Neurology, Mayo Clinic, Rochester, MN, USA; Department of Molecular Medicine, Mayo Clinic, Rochester, MN, USA

**Keywords:** microglia, antigen presentation, TMEV, viral infection, MHC class I, perivascular macrophage, CD8 T cell, atrophy

## Abstract

CD8 T cell infiltration of the central nervous system (CNS) is necessary for host protection but contributes to neuropathology. Antigen presenting cells (APCs) situated at CNS borders are thought to mediate T cell entry into the parenchyma during neuroinflammation. The identity of the CNS-resident APC that presents antigen via major histocompatibility complex (MHC) class I to CD8 T cells is unknown. Herein, we characterize MHC class I expression in the naïve and virally infected brain and identify microglia and macrophages (CNS-myeloid cells) as APCs that upregulate H-2K^b^ and H-2D^b^ upon infection. Conditional ablation of H-2K^b^ and H-2D^b^ from CNS-myeloid cells allowed us to determine that antigen presentation via H-2D^b^, but not H-2K^b^, was required for CNS immune infiltration during Theiler’s murine encephalomyelitis virus (TMEV) infection and drives brain atrophy as a consequence of infection. These results demonstrate that CNS-myeloid cells are key APCs mediating CD8 T cell brain infiltration.

## INTRODUCTION

T cells play essential roles in host protection from CNS infection; yet, infiltrating T cells induce potent immunopathology during neuroinflammation associated with infections and autoimmunity^1,2,3,4,5,6,7^. Recent studies suggest that dysregulated T cell brain infiltration may also contribute to pathology in neurodegenerative diseases^8,9,10^. Rediscovery of the glymphatic system in the CNS has allowed for improved understanding of how T cells are primed and activated against CNS-derived antigens^11, 12^. Conceivably, T cells encounter cognate antigen once it has drained to the deep cervical lymph node, where antigen presenting cells (APCs) prime naïve T cells^13, 14^. However, activated T cells are restimulated by local APCs to infiltrate across the blood brain barrier (BBB) into the parenchyma. This crucial process is not fully defined^15,16,17^.

CNS antigen presentation has been primarily studied in the experimental autoimmune encephalomyelitis (EAE) model of multiple sclerosis in which activated autoantigen-specific CD4 T cells migrate to the CNS, infiltrate the parenchyma, and mediate disease development^18^. Local APCs are crucial for CD4 T cell infiltration, and this role is primarily attributed to CD11c^+^ dendritic cells^19,20,21^. However, microglia and brain macrophages (CNS-myeloid cells) become fully competent APCs during CNS infection models, and the contribution of local APCs in potentiating CD8 T cell infiltration of the CNS remains undefined^22,23,24,25^. One viral model commonly used to assess immune infiltration of the brain is Theiler’s murine encephalomyelitis virus (TMEV), a neurotropic murine picornavirus. Immune infiltration mediates clearance of TMEV in C57BL/6 mice, but this results in cognitive deficits and brain atrophy^26,27,28^. However, mice deficient in CD8 T cells or lacking certain MHC class I haplotypes are unable to clear TMEV from the CNS, resulting in virus-induced demyelinating disease^29,30,31,32^. CD8 T cell activation, brain infiltration, and viral clearance are dependent on recognition of the immunodominant TMEV capsid protein-derived peptide VP2_121-130_ presented in the H-2D^b^ MHC class I molecule^30, 3113, 33^. The highly reproducible nature of this CD8 T cell response enables *in vivo* analysis of antigen presentation requirements for lymphocyte infiltration of the brain.

Here, we sought to define the role of local antigen presentation in CD8 T cell infiltration of the virally infected CNS. Activated CNS-myeloid cells upregulate MHC class I and localize near hippocampal vasculature during TMEV infection, a prime location to interact with CD8 T cells attempting to cross the BBB. Using transgenic C57BL/6 mice in which one of the two MHC class I molecules is deleted in CNS-myeloid cells, we uncovered differential requirements for H-2K^b^ and H-2D^b^ molecules in promoting immune infiltration of the brain. We further show that antigen presentation by CNS-myeloid cells promotes brain atrophy resulting from the CD8 T cell response against TMEV.

## MATERIALS AND METHODS

### Mice

C57BL/6J (B6; Stock No. 000664), B6.PL-Thy1/CyJ (Thy1.1; Stock No. 000406), B6.129P2(Cg)-Cx3cr1tm2.1(cre/ERT2)Litt/WganJ (CX3CR1creER, Stock No. 021160), and B6.129P2(Cg)-Cx3cr1^tm1Litt^/J (CX3CR1^GFP/GFP^; Stock No. 005582) were acquired from the Jackson Laboratories (Bar Harbor, ME). Following shipment, mice were acclimated for at least one week prior to use. CX3CR1-creER^T2^ x K^b fl/fl^ animals (CX3CR1^cre^/K^b^) and CX3CR1^creER^ x D^b fl/fl^ animals (CX3CR1^cre^/D^b^) were generated in house as described below. For experiments involving tamoxifen-mediated cre activation, male and female mice between 4 and 6 weeks of age received tamoxifen (or corn oil) prior to experimental use at 10-12 weeks of age. For all other experiments, male and female mice between 5-12 weeks of age were used. 7-14-week-old female or male mice were used for donors in bone marrow chimera experiments. Heterozygous CX3CR1^GFP/+^ were used for experiments. All mice were group housed under controlled temperature and humidity with a 12-h light/dark cycle. Mice were provided ad libitum access to food and water. All animal experiments were approved by and performed in accordance with the Mayo Clinic Institutional Animal Care and Use Committee and the National Institutes of Health guidelines.

### Generation of transgenic CX3CR1^creER^ x K^b fl/fl^ and CX3CR1^creER^ x D^b fl/fl^ mouse strains

Transgenic H-2K^b^ LoxP (H-2K^b fl/fl^) and H-2D^b^ LoxP (H-2D^b fl/fl^) mice were both generated by our group as previously described^13, 14^. In brief, LoxP sites were inserted into K^b^ and D^b^ transgenes which were previously cloned via site-directed mutagenesis. After K^b^ or D^b^ transgene insertion to C57BL/6J mice (Stock No. 000664) by the Mayo Clinic Transgenic Mouse Core (Rochester, MN), animals were backcrossed onto MHC class I deficient animals (H2K^b^D^b^ KO) resulting in mice in which the transgenic K^b^ or D^b^ was the only MHC class I molecule expressed. CX3CR1^creER^ (Stock No. 021160) animals were crossed to MHC class I deficient animals (H-2K^b^D^b^ KO) for at least three generations for the strain to be MHC class I deficient (The Jackson Laboratory, Bar Harbor, ME). MHC class I deficient CX3CR1^creER^ animals were then crossed to the K^b^ LoxP mouse to generate CX3CR1^creER^/K^b^ cKO animals, or the D^b^ LoxP mouse to generate CX3CR1^creER^/D^b^ cKO animals. Tail DNA screening was performed using polymerase chain reaction for cre using primer sequences recommended by Jackson Laboratory (Forward: AAG ACT CAC GTG GAC CTG CT; Mutant, Reverse: CGG TTA TTC AAC TTG CAC CA; WT, Reverse: AGG ATG TTG ACT TCC GAG TTG). Flow cytometry was used to confirm presence of the CX3CR1creER transgene via YFP reporter expression and class I deficiency using K^b^ and D^b^ surface expression. Mice were considered cKO if they were positive for the cre transgene by PCR and flow cytometry, and tamoxifen injection successfully mediated deletion of K^b^/D^b^ surface protein on CX3CR1-expressing cells.

### Tamoxifen administration

Tamoxifen (Sigma-Aldrich, St. Louis, MO) was administered in corn oil (Sigma-Aldrich, St. Louis, MO) at a concentration of 20mg/mL. Tamoxifen was dissolved in corn oil by shaking overnight at 37°C in the dark. Animals were administered 75mg/kg tamoxifen intraperitoneally at 4-6 weeks of age for five consecutive days using a 26 gauge 3/8” beveled needle. Post-tamoxifen recovery times of 10 days or 6 weeks were incorporated prior to use of mice in experiments. Experiments using vehicle control injections were performed in a similar manner using corn oil as vehicle.

### Acute TMEV and TMEV-OVA infection

The Daniel’s strain of TMEV was prepared as previously described^13^. TMEV-XhoI-OVA8 (TMEV-OVA) was generated and prepared by our group as previously described^34^. At 5-12 weeks of age, mice were anesthetized with 1-2% isoflurane and infected intracranially (i.c.) with 2 x 10^6^ PFU of the Daniel’s strain of TMEV, or 2 x 10^5^ PFU of TMEV-OVA. Virus was delivered to the right hemisphere of the brain in a final volume of 10 μL using an automatic 1 mL Hamilton syringe (Hamilton Company, Reno, NV). Mice were euthanized for flow cytometry or immunofluorescence at 0-, 5-, or 7-days post infection (dpi).

### PLX3397 administration

PLX3397 was synthesized by Plexxicon Inc. (Berkeley, CA) and formulated in AIN-76A standard chow by Research Diets Inc. (New Brunswick, NJ) at 290 mg/kg. AIN-76A standard chow alone was used as respective controls. Diets were provided to mice for two weeks ad libitum.

### BrdU administration

BrdU (BD Pharmingen, Cat#51-2420KC) was administered by intraperitoneal injection of a 100uL solution of 10mg/mL at day 6 post intracranial TMEV infection according to manufacturer’s instructions.

### Isolation of immune cells from secondary lymphoid organs

Spleens, cLNs, and thymi were harvested in 5 mL RPMI (RPMI 1640, Gibco) and gently homogenized between the frosted glass of two glass microscope slides. Samples were washed once at 400x*g* with RPMI in 15 mL conical tubes. Next, 1 mL of ACK lysis buffer (8.3 g ammonium chloride, 1 g potassium bicarbonate, and 37.2 mg EDTA) was added to the spleen samples for 1 minute to lyse erythrocytes. To quench the ACK reaction, 14 mL RPMI was added, and samples were washed once more prior to staining for flow cytometric analysis.

### Isolation of immune cells from whole brain

Immune cells were isolated from mouse brain as previously described^35^. Briefly, mice were deeply anesthetized with isoflurane and transcardially perfused with 30 mL of ice cold 1X PBS via intracardiac puncture. Whole brains were collected into 5 mL of ice cold RPMI and manually homogenized using a 7 mL glass Tenbroeck tissue grinder (Pyrex #7727-07). Homogenized brain samples were then filtered through a 70 µm filter (Falcon #352350) into a 30% Percoll gradient (Millipore Sigma, Darmstadt, Germany - #P4937) and centrifuged at 7840x*g*. The floating myelin debris layer was subsequently removed, and leukocytes were collected. Samples were washed twice with 1X PBS prior to staining for flow cytometric analysis.

### Flow cytometry

Cells were counted using a hemocytometer (Hausser Scientific) using trypan blue exclusion (Gibco) prior to being plated in a 96-well v-bottom plate. When applicable, samples were stained with 50 µL of a 1:50 dilution of D^b^:VP2_121-130_ APC-labeled tetramer or a 1:50 dilution of K^b^:OVA_257-264_ APC-labeled tetramer (NIH Tetramer Core Facility, Emory University). Tetramer staining was performed for 25 minutes in the dark at room temperature. Subsequently, samples were stained with the relevant combination of surface and intracellular antibodies in combination with Fc blocking antibody CD16/CD32 (BD Pharmingen, Cat. #553141). BV421 anti-MHCII(IA/IE) (BioLegend, Cat. #107632), Spark NIR 685 anti-CD45R/B220 (BioLegend, Cat. #103268), PE-CF594 anti-CD45 (BD Pharmingen, Cat #562420), PE anti-H-2Db (ThermoFisher, Cat. # A15443), PerCP anti-Ly6C (BioLegend, Cat. #128028), BB515 anti-CD11b (BD Pharmingen, Cat. #564454), APC/Fire750 anti-CD62L (BioLegend, Cat. #104450), PE-Cy7 anti-TCR*β* (Tonbo, Cat. #60-5961), Alexa Fluor 700 anti-H-2Kb (BioLegend, Cat. #116521), Pacific Blue anti-CX3CR1 (BioLegend, Cat. #149038), BV650 anti-CD44 (BD Pharmingen, Cat. #740455), BV510 anti-CD4 (BioLegend, Cat. #100449), BV570 anti-CD8*α*(BioLegend, Cat. #100740), BV605 anti-CD11c (BioLegend, Cat. #117333), BV711 anti-Ly6G (BioLegend, Cat. #127643), BV785 anti-F4/80 (BioLegend, Cat. #123141) antibodies were used at 1:100 dilution to stain cells from all tissues. Zombie NIR viability dye (BioLegend, Cat. #423105) was used at a 1:1000 dilution to stain dead cells. Samples were run on a BD LSRII flow cytometer equipped with FACSDiva software or a Cytek Aurora flow cytometer equipped with SpectroFlo software. Samples run on the LSR II were compensated with single stain controls, and samples run on the Cytek Aurora were unmixed with single stain reference controls.

### Processing of flow cytometry data

All samples were analyzed using FlowJo v10 (FlowJo LLC, Ashland, OR). Live, single, quality-controlled, and compensated/unmixed events covering 20 samples originating from naïve and infected B6 mice were equivalently downsampled to more than 1500 events per sample (https://www.flowjo.com/exchange/#/plugin/profile?id=25). We ran UMAP on the concatenated data, using 15 nearest neighbors (nn), a *min_dist* of 0.5, and Euclidean distance (https://arxiv.org/abs/1802.03426). Identified populations were confirmed by manual gating.

### Immunofluorescence and microscopy

Mice were deeply anesthetized with isoflurane and transcardially perfused with 30 mL of ice cold 1X PBS followed by 30 mL of ice cold 4% paraformaldehyde (PFA). For intravascular labeling experiments, mice were injected with 70 kDa Dextran conjugated to Texas Red (Invitrogen, Cat. #D1830) 7.5 minutes prior to anesthetization. Tissues were post-fixed overnight in 4% PFA at 4°C, then incubated 24h in 15% sucrose at 4°C, and finally incubated 24h in 30% sucrose at 4°C prior to embedding in Tissue-Tek OCT Compound (Sakura Finetek, Torrance, CA). Tissues were then sectioned at 20-um thickness by cryostat (Leica Biosystems, Wetzlar, Germany) onto positively charged glass slides. Slides were blocked for 1 hour in PBS containing 1% BSA, 10% normal goat serum, and 0.1% Triton-X 100 Sigma-Aldrich) prior to incubation overnight at 4°C with primary antibody diluted in block buffer: Rabbit anti-Iba1 (1:1000 019-19741, Wako, Osaka, Japan), Rabbit anti-NeuN (1:1000, Abcam, Cat. #104225). Slides were then washed with 1X PBS 3 times prior to incubation with fluorochrome-conjugated secondary antibody (goat anti-rabbit Alexa Fluor 647, ThermoFisher, Cat. #A-21245) for 1 hour at room temperature. Finally, slides were washed 5 times with 1X PBS prior to being mounted with VectaShield medium containing DAPI (Vector lab, Burlingame, CA).

Sections (greater than 3 sections per mouse) for Iba1+/CX3CR1+ cell density analysis or NeuN+ analysis were imaged with the Zeiss AxioObserver.Z1 structured illumination system (Carl Zeiss Microscopy GmbH, Jena, Germany) using a 40x objective. TIFF images were exported using Zen Blue software. For Iba1+ or CX3CR1+ cell morphology analysis (soma, Skeleton, and Sholl analysis), and vascular analysis, slides were imaged at room temperature using a Leica DM2500 (Wetzlar, Germany) equipped with a x63 oil immersion objective (confocal image: 521×521). 25-micrometer z-stacks were acquired with a step thickness of 1.01 um. Uncompressed TIFF images were exported from Leica Acquisition Suite software.

### Analysis and quantification of immunofluorescent images

All image analysis was performed using Fiji software (ImageJ, U. S. National Institutes of Health, Bethesda, Maryland, USA). Confocal z-stacks were smoothed and compressed into a single z-projection Confocal images were analyzed in Z-projected images (20 slices, maximum intensity projection). Cell numbers and density quantification was completed using the Fiji (ImageJ) Analyze Particles function after uniform thresholding. Vessel-associated CX3CR1+ cells were defined using methods described by Haruwaka et al^36^. Briefly, a vertical line was drawn from the presumed center of the soma of a CX3CR1+ cell (labeled by eGFP) to the outer surface of the blood vessel (labeled by dextran). The Multiplot function in Fiji (ImageJ) was utilized to calculate the relative fluorescence intensities of GFP and Texas Red along this line, then the standard deviation of fluorescent intensity was calculated for each channel. Finally, the distance between the point where the eGFP signal decreased to zero and the Texas Red signal increased from zero was assessed. Vessel-association was defined as a distance between green and red signals as below 1 μm, and the percent vessel-associated of total CX3CR1+ cells per ROI was calculated. For Sholl and Skeleton analysis, each maximum intensity projection Z-stack image was uniformly thresholded. The Sholl analysis plugin of ImageJ was applied^37^. The area under the curve generated by Sholl analysis per individual animal was calculated. CNS-myeloid cell soma size was measured using the segmented line tool in Fiji (ImageJ). Finally, a skeleton was created to assess CNS-myeloid cell morphology and branching using the AnalyzeSkeleton plugin^38^.

### Plaque assay

Infectious virus plaque assay was conducted using whole brain homogenates as previously described^39^. Briefly, whole brains were collected from deeply anesthetized mice (isoflurane) and perfused with 30 mL ice cold 1X PBS. Brains were weighed, then sonicated until fully homogenized. Homogenate was clarified by centrifugation at 1000x*g* for 20 minutes. L2 cells (ATCC CCL-149™) were grown in DMEM w L-glut (Gibco), 10% Heat-Inactivated Fetal Bovine Serum and 1% penicillin-streptomycin (Sigma) and plated onto 12-well plates at 1×10^6^ cells/well. The assay was performed once the cells reached confluency. Confluent cells were washed once with serum-free DMEM. Next, 10-fold dilutions of tissue homogenate were prepared in serum-free DMEM and 200 μL of each dilution was applied in triplicates onto cells. Plates were incubated at 37°C for 1 hour before 1 mL of a 0.8% agarose overlay was added. After 72 hours of incubation at 37°C, cells were fixed with EAF fixative (EtOH:HOAc:formaldehyde 6:2:1) for 1 hour prior to aspiration of agarose+fixative and staining with Crystal Violet solution (1% crystal violet in 20% EtOH). Plaques were counted by hand and PFU/g tissue was calculated.

### Bone marrow isolation and set up of chimeras

Femurs and tibias were isolated from donor mice. Spongey bones were cut off and bone marrow was flushed out under sterile conditions with a 21G needle. Cell suspensions were then lysed with ACK lysis buffer, washed twice at 400*xg*, and transferred intravenously into lethally irradiated recipient mice. Recipient mice received two doses of 450 gray irradiation four hours apart using a Shepherd’s CS 137 irradiator. Donor chimerism was assessed in peripheral blood at 6 weeks post-engraftment using flow cytometry.

### Novel object recognition testing

Novel object recognition testing was conducted as previously described^40^. In brief, mice were habituated to a 33 cm x 33 cm acrylic open field apparatus for 5 minutes per single mouse one day prior to the assay. The next day, mice were trained via exposure to two identical objects (red wooden blocks) arranged in opposite quadrants and 5 cm away from the walls. One mouse at a time was placed in the center of the open field with control objects and allowed to behave freely for 10 minutes. The mouse was then returned to the home cage for 1 hour prior to the novel object recognition testing session. Testing sessions consisted of a 5-minute exposure to one control object (the red wooden block) and one novel object (randomly assigned object differing in shape and texture made of Lego bricks), both placed in opposite quadrants and 5 cm away from the walls. All trials were conducted in a quiet, isolated procedure room under dim lighting. The open field apparatus and all objects were cleaned with 70% ethanol and dried in between each animal and each session. EthoVision XT software (Noldus Information Technology, Wageningen, the Netherlands) was used to record and count the number of investigations/sniffs of each object. The number of interrogations of the novel object as compared to the number of interrogations of the familiar object was calculated for the 5-minute test session (Discrimination Index = novel/familiar), and the number of interrogations of each control object were compared for the 10-minute training session (familiar/familiar). A mouse with equivalent exploratory behavior and learning/recognition memory would have a discrimination index of 1 during training and greater than 1 during testing, indicating more exploration of the novel object.

### Magnetic resonance imaging

Magnetic resonance images were acquired as previously described^27^. A Bruker Avance II 7 T vertical-bore small-animal animal system was used to conduct T2-weighted scans. Mice were anesthetized under isoflurane for the entirety of scanning. A rapid acquisition with relaxation enhancement (RARE) pulse sequence was used for T2 scanning with the following metrics: a TR of 1,500 ms, a TE of 65 ms, a RARE factor of 16, a FOV of 4.0 by 1.92, and a matrix of 200 by 96 by 96. Analyze 12.0 software was used to analyze MRI scans and generate 3D volumes (Biomedical Imaging Resource, Mayo Clinic).

### Statistical analysis and reproducibility

Sample sizes were chosen on the basis of standard power analysis conducted on historical experimental groups, with alpha = 0.05 and power of 0.8. Experimenters were blinded to the identity of experimental groups for non flow-cytometric experiments. GraphPad Prism 8.0.1 software (La Jolla, CA) was used to perform statistical analyses. Data are presented as mean +/- SD. All statistical tests were performed following verification of the assumptions on the distribution of data using Shapiro-Wilk normality test and D’Agostino-Pearson normality test, and non-parametric tests were used if assumptions were not met. Multiple independent groups were compared using one-way ANOVA with Tukey’s multiple comparison test, and two group comparisons were made using unpaired Student’s t-test or Welch’s student t-test, with alpha = 0.05 for significance. Specific tests used are detailed in figure legends.

### Data availability

All data and material used for this paper are available from the authors upon request.

## RESULTS

### CNS-myeloid cells are the primary immune cell that upregulates MHC class I during viral infection of the brain

In order to define APCs responding to intracranial TMEV infection, we isolated cells from the brains of healthy and TMEV infected C57BL/6 mice at 7 days post infection (dpi) and used high-dimensional flow cytometry accompanied with unbiased clustering (Fig. 1a). We chose 7 dpi as it is the peak of TMEV-specific CD8 T cell responses in C57BL/6 mice^41^. Using the Uniform Manifold Approximation and Projection for Dimension Reduction (UMAP) algorithm, we mapped live CD45^+^ populations onto a 2D space and identified 7 unique clusters of CD45^+^ cells, whose expression of various cell surface phenotypic markers broadly identified the population (Fig. 1b, Sup. Fig. 1a-b). We manually confirmed the identity of these populations based on known surface markers and assigned manual gates in subsequent experiments (Sup. Fig. 2).

**Figure 1.**
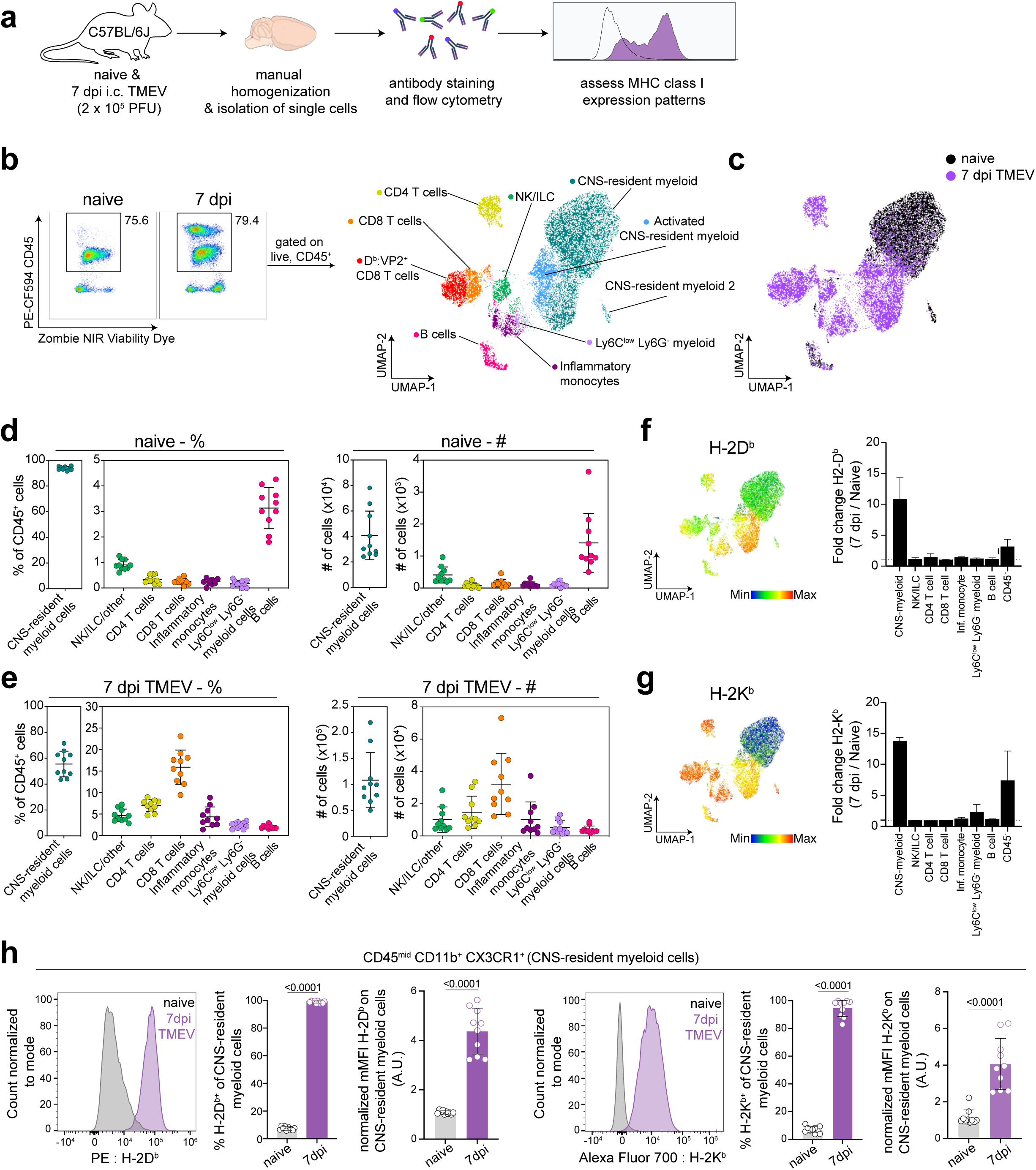
Analysis of brain immune compartment reveals CNS-myeloid cells upregulate MHC class I in response to CNS-viral infection. (a) Experimental procedure. Brains were isolated from perfused naïve and TMEV-infected mice at 7 days post infection (dpi) (n=10), dissociated using manual homogenization, and stained with fluorescently tagged antibodies for analysis of antigen presenting cells using high-dimensional flow cytometry. (b) Live CD45^+^ cells were downsampled and pooled for Uniform Manifold Approximation and Projection for Dimension Reduction (UMAP) analysis, identifying 7 immune cell clusters within naïve and TMEV-infected brains. (c) UMAP visualization color-coded by infection status. (d) The frequency and number of identified immune cells in the naïve brain. (e) The frequency and number of identified immune cell populations in the TMEV-infected brain at 7dpi. (f-g) UMAP visualization of CD45^+^ populations, colored by H-2D^b^ expression (f) and H-2K^b^ expression (g), and fold change of H-2D^b^ expression (f) and H-2K^b^ expression (g) at 7 dpi compared to naïve in CD45^+^ and CD45^-^ cell populations. (h) CNS-myeloid cells manually gated as CD45^mid^ CD11b^+^ CX3CR1^+^ were analyzed for H-2D^b^ and H-2K^b^ expression using flow cytometry. Representative histograms, frequency of H-2D^b^ and H-2K^b^ positive cells of the population, and normalized median fluorescence intensity (MFI) of H-2D^b^ and H-2K^b^ on CNS-resident myeloid cells. Data are representative of ≥ 3 independent experiments and presented as mean +/- SD, 2-tailed unpaired Student’s *t* test with ns p ≥ 0.05. dpi, days post infection.

Consistent with prior reports, we observed that the naive mouse brain harbors primarily CNS-resident myeloid cells (93.78% +/- 1.29) (Fig. 1b-c), which includes parenchymal microglia and perivascular macrophages. CD45^hi^ B220^+^ B cells were the second most abundant immune cells in the naïve brain (3.134 % +/- 0.77) (Fig. 1d). The remaining CD45^+^ cells consisted of CD8 T cells (CD45^hi^ TCRβ^+^ CD8α^+^, 0.26% +/- 0.11), CD4 T cells (CD45^hi^ TCRβ^+^ CD4^+^, 0.34% +/- 0.15), inflammatory monocytes (CD45^hi^ CD11b^+^ Ly6C^hi^ Ly6G^-^, 0.24% +/- 0.11), Ly6C^low^ infiltrating myeloid cells (CD45^hi^ CD11b^+^ Ly6C^low^ Ly6G^-^, 0.16% +/- 0.12), and ILC/NK/other (CD45^hi^ CD11b^-^ TCRb^-^B220^-^, 0.907 +/- 0.19) (Fig. 1d). In comparison, at 7 days post TMEV infection, hematopoietic cells infiltrate the brain, resulting in reduced proportions of CNS-myeloid cells (55.53% +/- 9.39 of the total CD45^+^ population, Fig. 1e). CD8 T cells predominated the infiltrate, making up 15.91% +/- 3.77 of the CD45^+^ cells, followed by 6.96% +/- 1.30 CD4 T cells, 4.361% +/- 2.18 inflammatory monocytes, 2.40% +/- 0.68 Ly6C^low^ infiltrating myeloid cells, 2.00% +/- 0.36 B cells, and 4.68% +/- 1.40 NK/ILC/other (Fig. 1e). We did not detect neutrophils in our analyses. There were no significant differences in cell proportions or numbers when male and female mice were compared (Sup. Fig. 1a).

We then assessed MHC class I expression by these populations. In the C57BL/6 mouse, two MHC class I molecules are expressed: H-2K^b^ and H-2D^b^. We found that CD45^hi^ populations expressed both K^b^ and D^b^ during the steady state, though the median fluorescence intensity (MFI) varied between populations indicating differences in expression levels (Fig. 1f-g). CNS-myeloid cell expression of K^b^ or D^b^ was undetectable at steady state, but both class I molecules were markedly upregulated during TMEV infection. UMAP visualization colored by H-2D^b^ expression indicates that the highest relative H-2D^b^ expressing cells were CNS-myeloid cells and infiltrating myeloid cells at 7 dpi (Fig. 1f), while CNS-myeloid cells expressed relatively less H-2K^b^ compared to other CD45^+^ cells at 7 dpi (Fig. 1g). The robust increase in MHC class I expression during TMEV infection was unique to CNS-myeloid cells, as demonstrated by fold change when all CD45^+^ cell types were accounted for (Fig. 1f-g). CD45^-^ cells upregulated K^b^ and D^b^ but to a lesser extent than CNS-myeloid cells (Fig. 1f-g). Overall, CNS-myeloid cells had the largest increase in MHC class I expression resulting from TMEV infection (Fig. 1h).

In addition to upregulation of MHC class I, CNS-myeloid cells upregulated MHC class II (IA/IE) at 7 days post TMEV infection (Sup. Fig. 3a). Previous studies contend that MHC class II upregulation was limited in experimental autoimmune encephalomyelitis (EAE) models^20^. Nonetheless, about 45.81% +/- 11.28 of CNS-myeloid cells upregulated MHC class II at 7 days post TMEV infection, indicating that CNS-myeloid cells can express MHC class II (Sup. Fig. 3a). Together, these results demonstrate that CNS-myeloid cells, the most abundant immune cells in the naïve and infected brain, upregulate both MHC class I and MHC class II during acute TMEV infection. The striking upregulation of MHC class I expression by CNS-myeloid cells suggests that CNS-myeloid cells have the potential to present antigen to infiltrating CD8 T cells during acute TMEV infection.

### CNS-myeloid cells are activated and associate with hippocampal vasculature during acute TMEV infection

Activation of CNS-myeloid cells is a common response to injury, neuroinflammation, and neurodegeneration^42, 43^. We observed that CNS-myeloid cells markedly upregulated MHC class I during TMEV infection, leading us to wonder whether this was a facet of CNS-myeloid cell activation during infection. Accordingly, we assessed CNS-myeloid cell activation in the hippocampus at 7 days post TMEV infection, as the hippocampus has been reported to be a primary site of infection in C57BL/6 mice after intracranial inoculation^33^. TMEV infection induced an increase in the density of hippocampal CNS-myeloid cells, indicating CNS-myeloid cell activation (Fig. 2a-b). We noticed CNS-myeloid cell clustering around vasculature (Fig. 2a, inset), and measured increased association of CNS-myeloid cells with vasculature during TMEV infection (Fig. 2c-d). Thus, CNS-myeloid cells are positioned for interactions with CD8 T cells infiltrating the hippocampus during acute TMEV infection. Additional morphologic changes in CNS-myeloid cells were measured in TMEV infected mice, including changes from a resting “ramified” morphology to a reactive “bushy” morphology (Fig. 2e), increased soma size (Fig. 2f), and reduced microglial process branching and complexity during infection (Fig. 2g). Flow cytometric analyses permitted us to measure additional activation metrics in CNS-myeloid cells during TMEV infection. Infection increased the frequency of CX3CR1^mid^ side-scatter (SSC)^low^ CNS-myeloid cells that were absent under steady state conditions (Sup. Fig. 3b-d). SSC changes were consistent with CNS-myeloid cell morphological alterations observed in the hippocampus (Fig. 2), and reduced CX3CR1 expression is related to the activated “damage associated microglia” (DAM) phenotype seen in neurodegenerative diseases^44^. The frequency of K^b^, D^b^, and MHC class II expressing cells was similar between CNS-myeloid cell stratifications, though differences in MFI suggest differential levels of surface protein expression indicating contrasting activation levels between CNS-myeloid cells during TMEV infection (Sup. Fig. 3e). Together, our data indicate that TMEV infection induces activation of CNS-myeloid cells that includes association with hippocampal vasculature and upregulation of MHC molecules.

**Figure 2.**
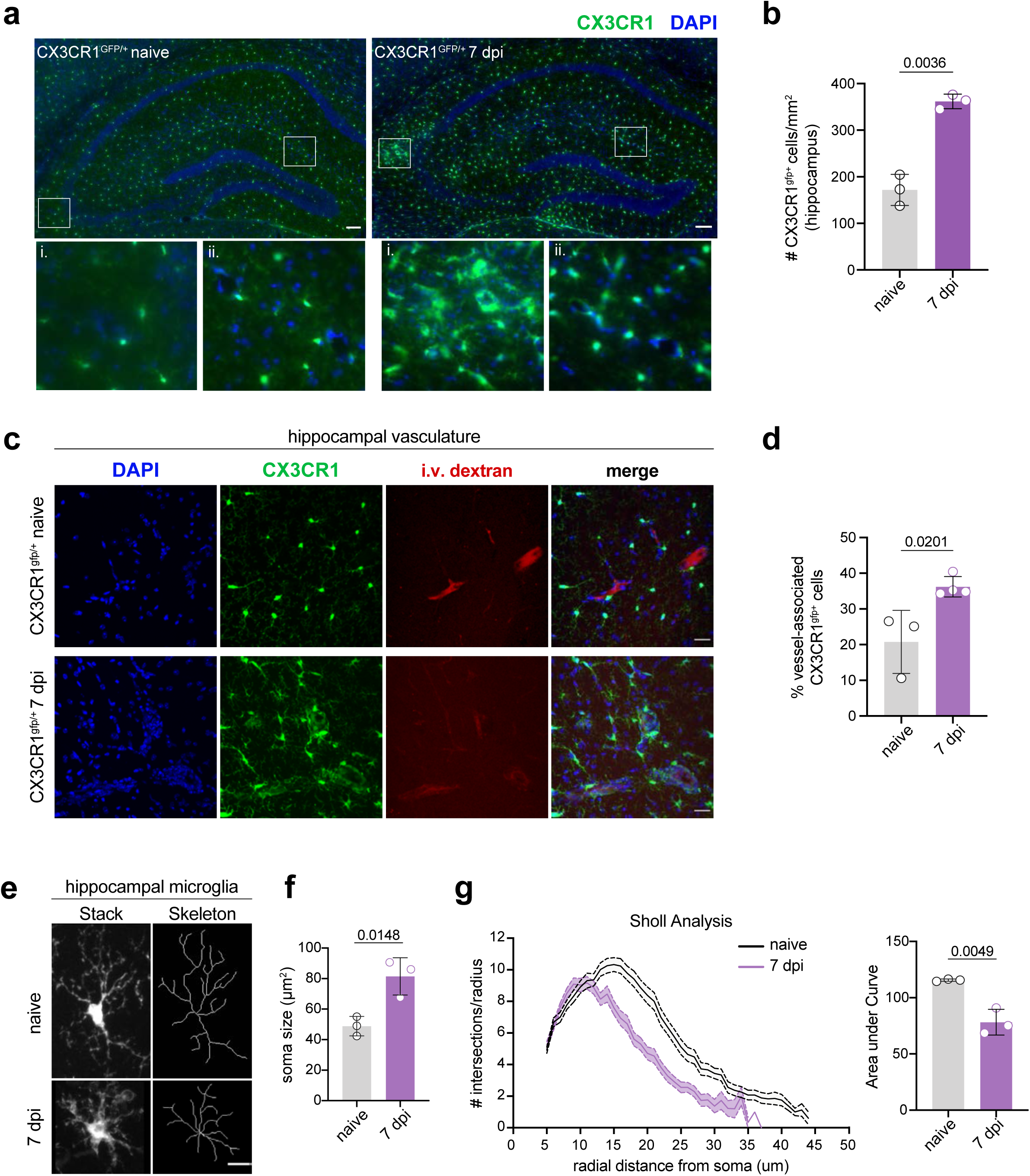
TMEV infection induces CNS-myeloid cell activation and vessel association in the hippocampus. (a) CX3CR1^+^ CNS-myeloid cells (green) in representative hippocampal images from uninfected and TMEV infected (7 dpi) CX3CR1^gfp/+^ mice. Zoomed insets demonstrate areas in which CX3CR1^+^ cells cluster around vasculature. *n* = 3 mice (3 sections/mouse). Scale bars: 100 μm. (b) Density of hippocampal CX3CR1^+^ cells in naïve and TMEV infected mice. (c) Representative images of DAPI (nuclei, blue), CX3CR1 (CNS-myeloid, green), and intravascularly injected dextran (vasculature, red) in the hippocampus of naïve and TMEV-infected mice. *n* = 3 mice (6 sections/mouse). Scale bars: 25 μm. (d) Quantification of the proportion of CX3CR1^+^ cells in contact with hippocampal vasculature in naïve and infected conditions. (e) Representative images of hippocampal microglial morphology (confocal and transformed skeletal) in uninfected and TMEV infected mice (7 dpi). *n* = 3 mice (3 sections/mouse). Scale bars: 10 μm. (f) Quantification of hippocampal microglia soma size at 7 dpi TMEV compared to naïve. (g) Sholl analysis of microglia at 7 days post TMEV infection or during steady state. (25 microglia/section, 2 sections/mouse). Area under the curve of the Sholl analysis is used to demonstrate quantification of microglial branching during infection. Data are representative of ≥ 2 independent experiments and presented as mean +/- SD, 2-tailed unpaired Student’s *t* test with ns p ≥ 0.05.

### CSF1R inhibition alters peripheral myeloid cell populations and CD8 T cell priming

To assess whether CNS-myeloid cells were critical APCs for CD8 T cell infiltration of the CNS, we sought to deplete CNS-myeloid cells prior to TMEV infection. Colony stimulating factor 1 receptor (CSF1R) signaling is essential for the development and survival of myeloid cells, and CSF1R inhibitors have been utilized to deplete murine microglia in experimental settings^45^. We depleted CNS-myeloid cells by providing mice with the CSF1 inhibitor PLX3397 or control chow for 2 weeks. We then infected these mice with TMEV upon the start of week 2 (Sup. Fig. 4a). PLX3397 treated mice were protected from acute weight loss as a result of TMEV infection (Sup. Fig. 4b) but began to succumb to infection at 5 dpi. PLX3397 depleted CNS-myeloid cells as previously reported (Sup. Fig. 4c). However, we found that PLX3397 treatment altered the total number of cells isolated from the spleen (Sup. Fig. 4d) and the proportion and number of CD11c+ cells and MHC II+ cells, implying peripheral T cell priming by dendritic cells would be impaired (Sup. Fig. 4d). Indeed, virus-specific CD8 T cells primed in the spleen were reduced upon PLX3397 treatment (Sup. Fig. 4d). Taken together, we determined that PLX3397 was not specific enough for our experimental question due to off-target effects on peripheral immunity in addition to CNS-myeloid cell ablation.

### Development of a CNS-myeloid cell MHC class I deletion model

Our approach using CSF1R inhibition to reduce CNS-myeloid cells unintentionally disrupted peripheral immunity. Therefore, we developed a more CNS-specific approach to dissect the role of CNS-myeloid cell antigen presentation using conditional ablation of H-2K^b^ or H-2D^b^ expression. CX3CR1^creER^ mice on a MHC class I deficient background were crossed to either H-2K^b fl/fl^ or H-2D^b fl/fl^ transgenic mice that were otherwise MHC class I deficient^13, 14^. This resulted in two strains of mice carrying the CX3CR1^creER^ transgene, with either a floxed H-2K^b^ gene (CX3CR1^cre^/K^b^) or H-2D^b^ gene (CX3CR1^cre^/D^b^) but no other MHC class I molecules. We specifically deleted H-2K^b^ or H-2D^b^ in CNS-myeloid cells by taking advantage of the long-lived, self-renewing nature of these cells in comparison to peripheral CX3CR1-expressing cells. To this end, we allowed 6-weeks to pass after tamoxifen treatment while peripheral CX3CR1-expressing cells were restored by H-2K^b^ or H-2D^b^ sufficient hematopoietic precursors, and CNS-myeloid cells remained H-2K^b^ or H-2D^b^ deficient. We then used flow cytometry to assess MHC class I deletion efficiency (Fig. 3a-b).

**Figure 3.**
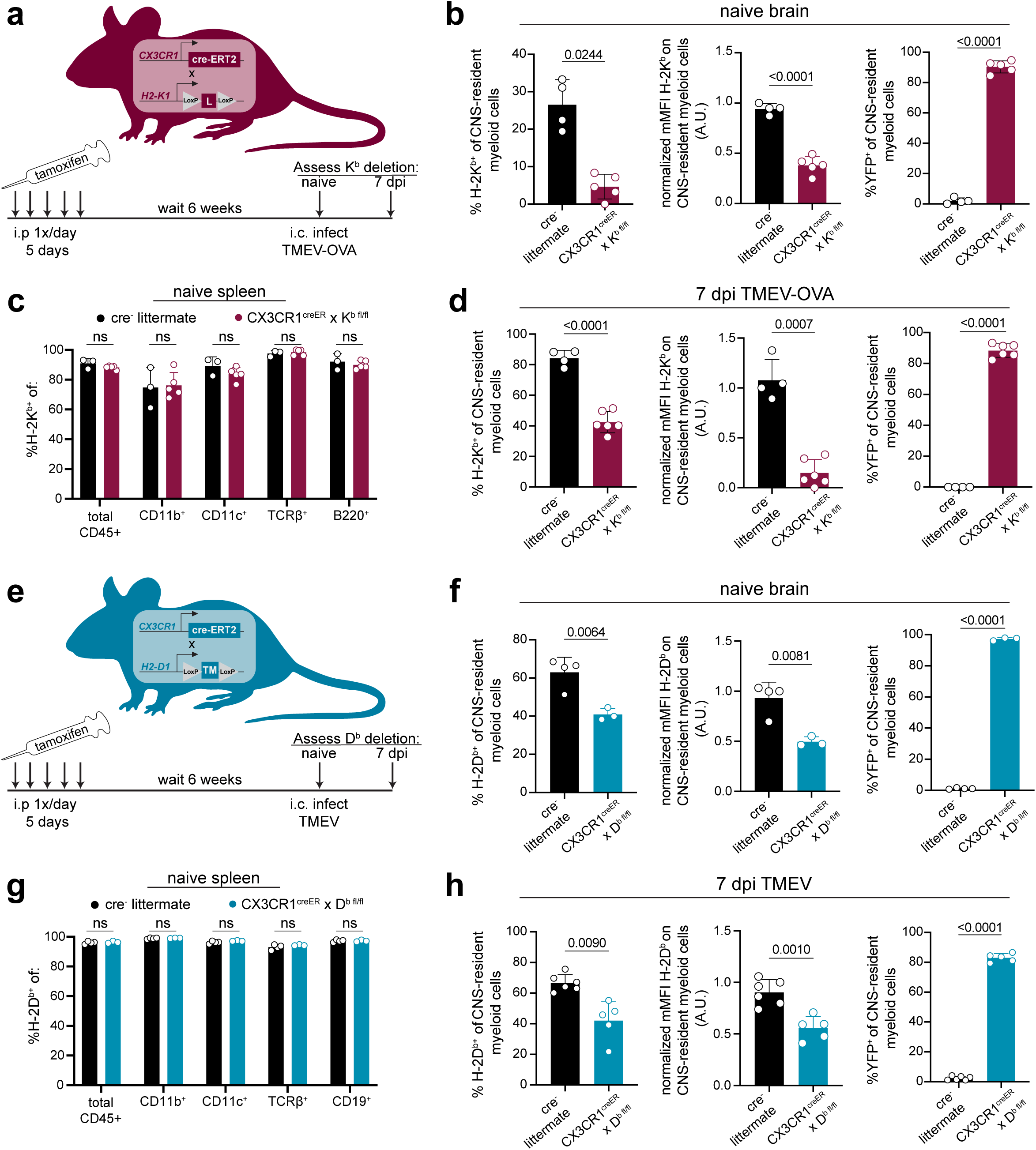
Development of a model to conditionally delete MHC class I in CNS-myeloid cells. (a, e) MHC class I deficient *CX3CR1^creER^* mice were crossed to transgenic H-2Kb ^fl/fl^ mice^14^ (a) or H-2Db ^fl/fl^ mice^13^ (e). Tamoxifen was used to induce MHC class I deletion. After 6 weeks, K^b^/D^b^ deletion was assessed using flow cytometry. (a-d) H-2K^b^ expression by CNS-myeloid cells was quantified by measuring percent positive and normalized median fluorescence intensity (MFI) in naïve CX3CR1^cre^/K^b^ mice and cre^-^ controls. YFP expression was measured to assess penetrance of CX3CR1^creER^ transgene. (c) H-2K^b^ expression was assessed in spleens of CX3CR1^cre^/K^b^ mice and cre^-^ controls. (d) H-2K^b^ expression by CNS-myeloid cells was quantified at 7 days post infection with TMEV expressing OVA_257-264_. (f) H-2D^b^ expression by CNS-myeloid cells was quantified by measuring percent positive and normalized median fluorescence intensity (MFI) in naïve CX3CR1^cre^/D^b^ mice and cre^-^ controls. YFP expression was measured to assess penetrance of CX3CR1^creER^ transgene. (g) H-2D^b^ expression was assessed in spleens of CX3CR1^cre^/D^b^ mice and cre^-^ controls. (h) H-2D^b^ expression by CNS-myeloid cells was quantified at 7 days post infection with TMEV. For all experiments, CNS-myeloid cells are live, single cells, expressing CD45^mid^CD11b^+^CX3CR1^+^. Data are shown as individual mice with mean from one independent experiment (n=3-6) of ≥ 3 experimental replicates. Error bars represent standard deviation. Two tailed Welch’s *t* test was used to assess statistical significance with ns p ≥ 0.05.

The CX3CR1^cre^/K^b^ system led to the robust deletion of H-2K^b^ in CNS-myeloid cells at baseline and during infection with TMEV expressing the model antigen ovalbumin (OVA_257-264_), which is presented in the K^b^ MHC class I molecule (Fig. 3b,d) ^34^. High YFP expression indicated that the majority of CNS-myeloid cells were targeted by this transgenic creER approach during naïve and infected conditions (Fig. 3b). Expression of H-2K^b^ was efficiently restored on peripheral immune cells through hematopoiesis, as there were no differences in the frequency of H-2K^b+^ CD45^+^ cells in the spleens of cre^+^ mice compared to cre^-^ controls (Fig. 3c). CX3CR1^cre^/D^b^ mice were similarly evaluated (Fig. 3e-h). CNS-myeloid cells from CX3CR1^cre^/D^b^ mice expressed significantly less H- 2D^b^ than cre^-^ animals during naive conditions and during infection with TMEV (Fig. 3f,h). We observed appropriate surface expression of H-2D^b^ in splenic immune cells (Fig. 3g). These results demonstrate that CX3CR1^cre^/K^b^ mice and CX3CR1^cre^/D^b^ have sufficient deletion of K^b^ or D^b^ expression by CNS-myeloid cells to allow assessment of antigen presentation by this cell type.

It has been reported that cre^ERT2^ lines such as the CX3CR1^creER^ used in this study have the potential to exhibit spontaneous, tamoxifen-independent excision of LoxP sites^46^, so we assessed the extent to which this occurred in our conditional knockout mice. We determined that CX3CR1^cre^/D^b^ mice treated with vehicle injections exhibited D^b^ expression comparable to cre^-^ controls (Sup. Fig. 5a-c). Accordingly, cre^-^ littermate controls were appropriate for the remainder of our studies. Altogether, we concluded that CX3CR1^cre^/K^b^ and CX3CR1^cre^/D^b^ mice are robust systems to specifically delete CNS-myeloid cell H-2K^b^ or H-2D^b^ during neuroinflammation, providing a valuable tool for studying CNS-myeloid cell antigen presentation without unintended consequences on peripheral immunity.

### CX3CR1-dependent H-2K^b^ or H-2D^b^ deletion does not impact CNS-myeloid cell homeostasis or T cell development

We next sought to investigate if there were any cell-intrinsic consequences of loss of MHC class I on CNS-myeloid cells. We determined that the numbers of CNS-myeloid cells isolated from the brains of CX3CR1^cre^/K^b^ and CX3CR1^cre^/D^b^ animals were comparable to cre^-^ controls at baseline and during infection (Sup. Fig. 6a-d). These data indicate that a loss of MHC class I expression on CNS-myeloid cells does not impact the survival of CNS-myeloid cells in the naïve or inflamed CNS. We then assessed whether loss of MHC class I had an impact on CNS-myeloid cell activation by performing Iba1 immunostaining in naïve- and TMEV-infected CX3CR1^cre^/D^b^ hippocampi. We determined that CNS-myeloid cell activation in response to TMEV infection was not impacted by CNS-myeloid cell loss of H-2D^b^ expression (Sup. Fig. 6e-g).

Following our analysis of the cell-intrinsic impact of MHC class I deletion, we sought to determine if conditional deletion of MHC class I on CX3CR1-expressing cells impacted T cell development. Using flow cytometry, we determined that thymocyte populations were unaffected by loss of K^b^ or D^b^ by CX3CR1-expressing cells (Sup. Fig. 7a-c, e-g). Further, splenic CD4 and CD8 T cell frequencies were similar in CX3CR1^cre^/K^b^ and CX3CR1^cre^/D^b^ mice compared to controls (Sup. Fig. 7d,h). Together, these data indicate that loss of MHC class I molecules on CX3CR1-expressing cells does not impact CNS-myeloid cell survival and activation, nor T cell development, providing a precise tool to dissect CNS-myeloid cell antigen presentation.

### H-2K^b^ expression by CNS-myeloid cells is dispensable for CD8 T cell responses to a model antigen

After confirming that CX3CR1^cre^/K^b^ animals served as a specific model of K^b^ deletion from CNS-myeloid cells, we sought to test whether CNS-myeloid cell H-2K^b^ is required for brain infiltration of CD8 T cells. Infection of C57BL/6 mice by TMEV engineered to express the model antigen OVA_257-264_ (TMEV-OVA) generates CD8 T cell responses against viral antigens and OVA^34^. We intracranially infected CX3CR1^cre^/K^b^ animals and controls with TMEV-OVA (consistent with the experimental design detailed in Fig. 3a) and assessed CD8 T cell responses in the spleen, cervical lymph node (cLN), and brains of these mice. The cLN is the putative site of T cell priming against CNS-drained antigens^12, 13^, and the spleen reflects systemic CD8 T cell responses against virus. The frequencies of CD8 T cells and OVA-specific CD8 T cells in the spleens and cLNs of CX3CR1^cre^/K^b^ and cre^-^ controls were comparable (Fig. 4a-b, d). Further, the frequencies of naïve (CD44-CD62L+), effector (CD44+CD62L-), and central memory (CD44+CD62L+) CD8 T cells in the spleens of CX3CR1^cre^/K^b^ mice did not differ from cre^-^ controls (Fig. 4c). These data indicate that T cell priming occurs normally in CX3CR1^cre^/K^b^ mice, which was expected, given K^b^ expression in the periphery was unaltered. We next sought to determine the extent of immune infiltration into the TMEV-OVA infected brain when CNS-myeloid cell K^b^ expression was disrupted. We found no differences in CD8 T cell (Fig. 4e-f), OVA-specific CD8 T cell (Fig. 4g-h), CD4 T cell (Fig. 4i), or inflammatory monocyte (Fig. 4j) infiltration of the brain, indicating H-2K^b^ MHC class I expression by CNS-myeloid cells is dispensable for neuroinflammation during TMEV-OVA infection of the brain.

**Figure 4.**
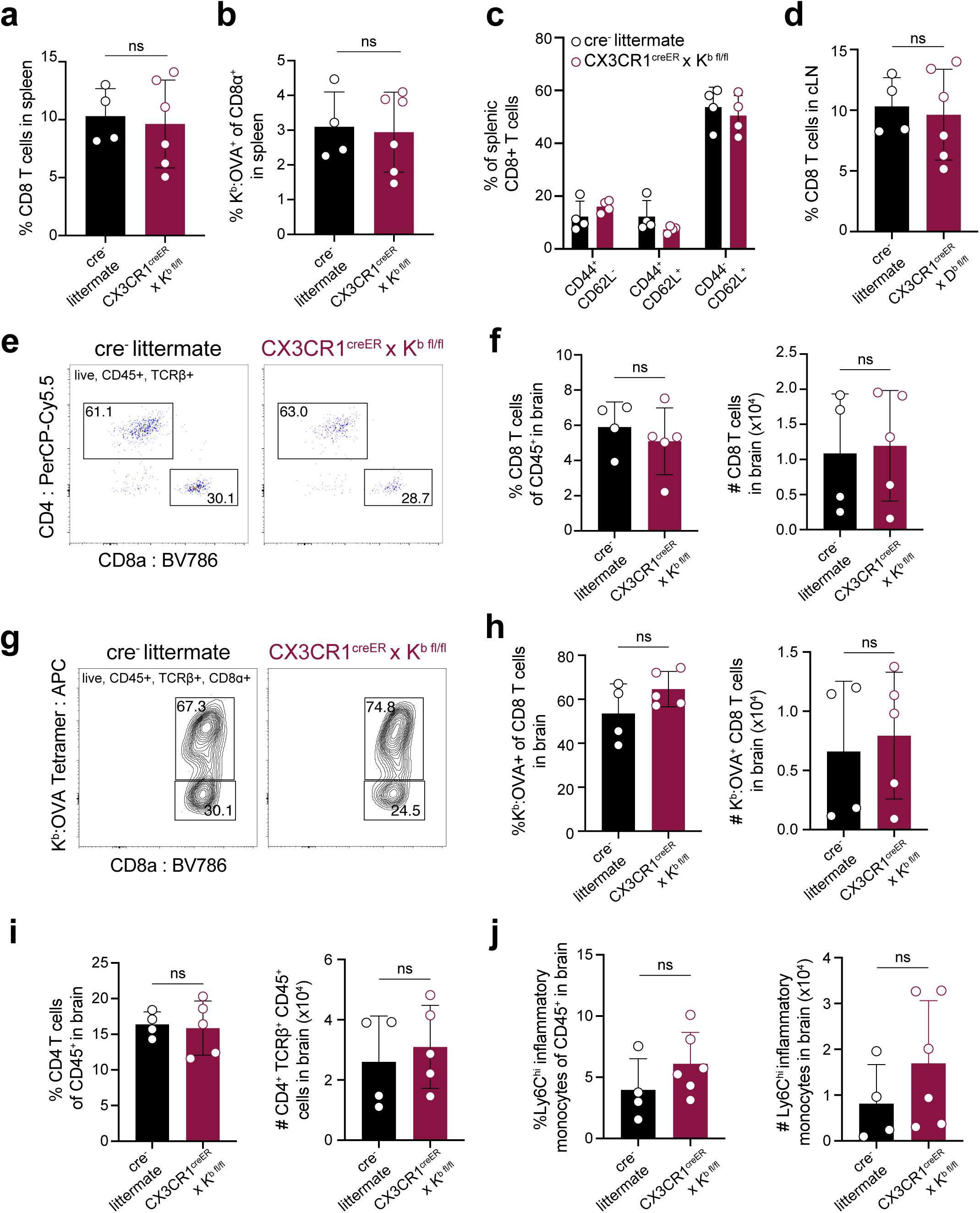
Deletion of H-2K^b^ in CNS-myeloid cells does not impact CD8 T cell responses to the K^b^ restricted antigen OVA_257-264_ during CNS-viral infection. Experimental strategy is identical to Figure 3a, in which tamoxifen administration in CX3CR1^cre^/K^b^ mice and cre^-^ controls is followed by a 6-week reconstitution period before intracranial (i.c.) infection with TMEV-OVA_257- 264_. Spleens, lymph nodes, and brains were isolated and analyzed by flow cytometry at 7 days post infection to assess antiviral responses. (a-b) Quantification of CD8^+^ T cells (a) and K^b^:OVA_257- 264_ CD8 T cells (b) in the spleens of infected mice. (c) Proportion of naïve (CD44 CD62L), effector (CD44^+^CD62L^-^), and central memory (CD44^+^CD62L^+^) CD8 T cells in the spleens of infected mice. (d) Quantification of CD8^+^ T cells in the CNS-draining deep cervical lymph node (dCLN) of infected animals. (e) Representative flow cytometry plots, and (f) quantification of total CD8 T cells infiltrating the brain at 7 days post infection in CX3CR1^cre^/K^b^ mice compared to controls. (g) Representative flow cytometry plots, and (h) quantification of total K^b^:OVA^+^ CD8 T cells infiltrating the brain at 7 days post infection. (i) Quantification of CD4 T cells recovered from the brains of cre^-^ and CX3CR1^cre^/K^b^ mice during infection. (j) Inflammatory monocytes recovered from the brains of infected cre^-^ and CX3CR1^cre^/K^b^ mice are quantified. Data are shown as individual mice with mean from one independent experiment (n=3-6) of ≥ 3 experimental replicates. Error bars represent standard deviation. Two tailed Welch’s *t* test was used to assess statistical significance with ns p ≥ 0.05.

### H-2D^b^ expression by CNS-myeloid cells is pivotal for neuroinflammatory responses to TMEV

Because we observed that CNS-myeloid cell expression of K^b^ did not impact immune infiltration of the brain during infection with TMEV-OVA (Fig. 4), we next sought to investigate CNS-myeloid cell antigen presentation via the D^b^ molecule. H-2D^b^, but not H-2K^b^, has been mapped to resistance from chronic TMEV infection^47,48,49^. This is attributed to viral clearance by CD8 T cell responses against the immunodominant viral VP2_121-130_ peptide antigen presented in context of the D^b^ class I molecule^30^. Therefore, we asked whether CNS-myeloid cell H-2D^b^ expression is required for brain infiltration of D^b^:VP2_121-130_ epitope specific CD8 T cells during acute TMEV infection. We intracranially infected CX3CR1^cre^/D^b^ animals and controls with TMEV (same experimental paradigm as Fig. 3e) and assessed CD8 T cell responses in the spleen, cLN, and brains of these mice. The frequencies of CD8 T cells, and D^b^:VP2 epitope specific CD8 T cells, in the spleens and cLNs of CX3CR1^cre^/D^b^ and control mice were comparable demonstrating that T cell priming is unaltered in CX3CR1^cre^/D^b^ mice (Fig. 5a-b, d). The frequencies of naïve (CD44- CD62L+), effector (CD44+CD62L-), and central memory (CD44+CD62L+) CD8 T cells in the spleens of CX3CR1^cre^/D^b^ and control mice also did not differ between genotypes (Fig. 5c). These data indicate that CD8 T cell priming occurs normally in CX3CR1^cre^/D^b^ mice, which is consistent with the observation that peripheral D^b^ expression was equivalent (Fig. 3g).

**Figure 5.**
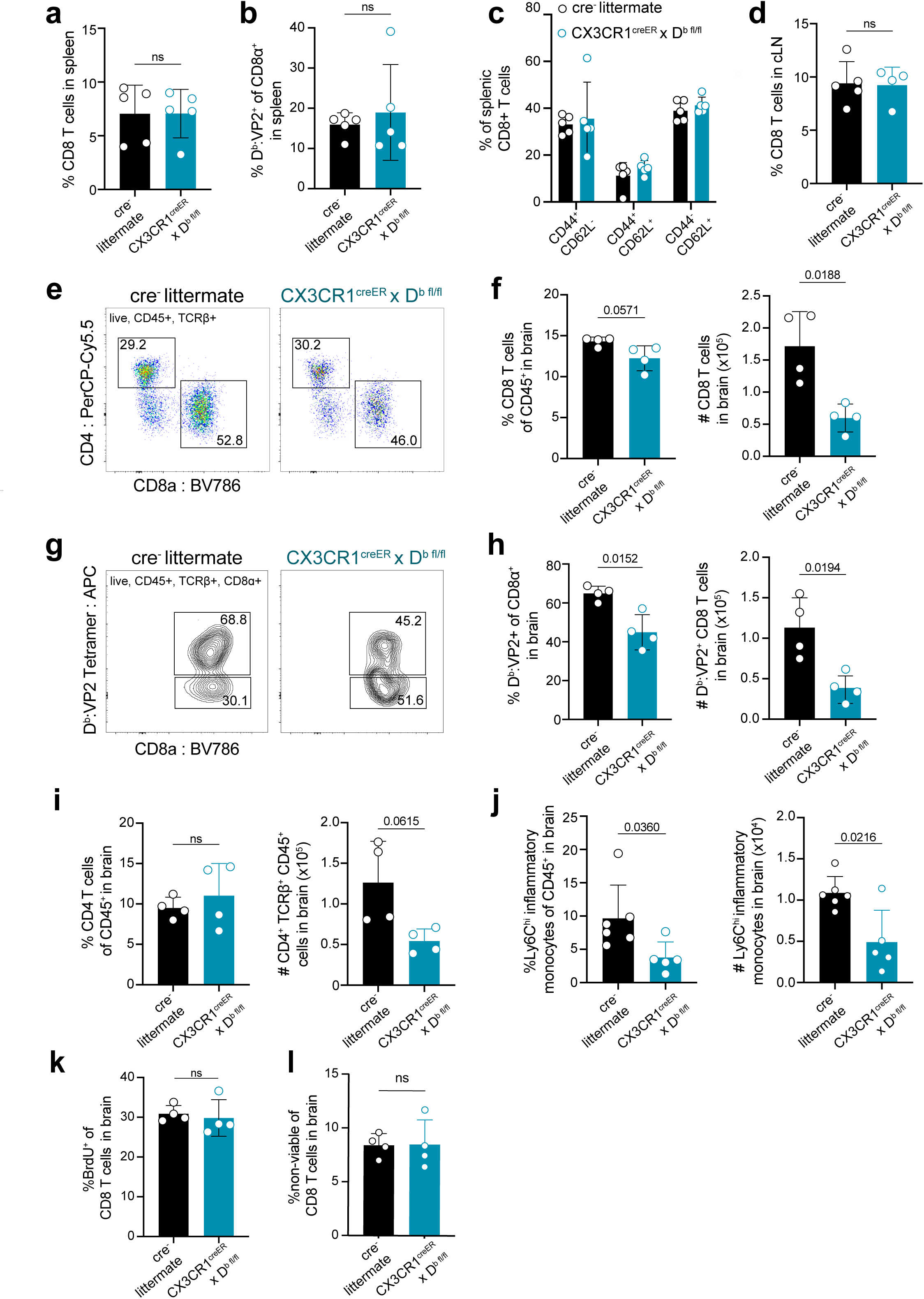
Deletion of H-2D^b^ in CNS-myeloid cells reduces neuroinflammatory responses to the D^b^ restricted viral antigen VP2_121-130_ during TMEV infection of the CNS. Experimental strategy is identical to Figure 3e, in which tamoxifen administration in CX3CR1^cre^/D^b^ mice and cre^-^ controls is followed by a 6-week reconstitution period before i.c. infection with TMEV. Spleens, lymph nodes, and brains were analyzed by flow cytometry at 7 days post infection to assess antiviral responses. (a-b) Quantification of CD8^+^ T cells (a) and D^b^:VP2^+^ CD8 T cells (b) in the spleens of infected mice. (c) Proportion of naïve (CD44^-^CD62L^+^), effector (CD44^+^CD62L^-^), and central memory (CD44^+^CD62L^+^) CD8 T cells in the spleens of infected mice. (d) Quantification of CD8^+^ T cells in the dCLN of infected animals. (e) Representative flow cytometry plots, and (f) quantification of CD8 T cells infiltrating the brain at 7 dpi in CX3CR1^cre^/D^b^ mice compared to controls. (g) Representative flow cytometry plots, and (h) quantification of total D^b^:VP2^+^ CD8 T cells infiltrating the brain at 7 dpi. (i) Quantification of CD4 T cells recovered from the brains of mice during infection. (j) Inflammatory monocytes recovered from the brains of infected mice are quantified. (k) BrdU was administered intraperitoneally at day 6 post infection and BrdU incorporation into brain-infiltrating CD8 T cells was quantified. (l) Viability dye incorporation was measured in CD8 T cells isolated from the brains of infected mice. Data are shown as individual mice with mean from one independent experiment (n=3-6) of ≥ 3 experimental replicates. Error bars represent standard deviation. Two tailed Welch’s *t* test was used to assess statistical significance with ns p ≥ 0.05.

In contrast to the equivalent peripheral T cell responses, we recovered reduced brain infiltrating cells in CX3CR1^cre^/D^b^ mice at 7 dpi with TMEV suggesting that immune infiltration of the brain was affected (Fig. 5). CX3CR1^cre^/D^b^ mice had a diminished number of total infiltrating CD8 T cells in the brain, as well as reduced virus specific (D^b^:VP2_121-130_ epitope specific) CD8 T cells infiltrating the brain (Fig. 5 e-h). We also noted a reduction in the frequency of inflammatory monocytes in the brains of CX3CR1^cre^/D^b^ animals as compared to controls (Fig. 5j). There were no other differences in immune infiltrates observed in CX3CR1^cre^/D^b^ animals, harboring comparable CD4 T cells responses (Fig. 5i), indicating the reduced immune infiltration of the brain was attributable to CNS-myeloid cell D^b^ loss.

To determine whether the reduced number of infiltrating T cells in CX3CR1^cre^/D^b^ mice was due to decreased T cell proliferation, infected CX3CR1^cre^/D^b^ and cre^-^ animals were given bromodeoxyuridine (BrdU) intraperitoneally at day 6 post infection and assessed at day 7 post infection. BrdU incorporation in CD8 T cells revealed no changes in T cell proliferation in CX3CR1^cre^/D^b^ mice, indicating that the reduced CD8 T cell response in the brain was not due to proliferation defects (Fig. 5k). Given similar cellular proliferation, we asked whether CD8 T cells infiltrating the brain experienced increased cell death, but uncovered no differences in T cell viability in CX3CR1^cre^/D^b^ mice compared to controls (Fig. 5l). Thus, CNS-myeloid cell antigen presentation has a critical role for CD8 T cell and inflammatory monocyte infiltration when these cells are presenting the immunodominant antigen during TMEV infection, independent of T cell proliferation or viability.

### CNS-myeloid cell antigen presentation impacts TMEV specific CD8 T cell responses at the point of brain infiltration

Our approach targeting CX3CR1-expressing cells to conditionally delete H-2D^b^ is designed to impact only long-lived CNS-myeloid cells. However, to further rule out the contribution of peripheral APCs towards CD8 T cell infiltration, we sought to confirm our findings using a bone marrow chimeric approach. Cre^-^ littermates and CX3CR1^cre^/D^b^ animals were lethally irradiated and provided with wild type bone marrow, generating mice in which the CX3CR1^creER^ transgene is only present in the CNS immune compartment while hematopoietic cells are wild type (Fig. 6a). In chimeric CX3CR1^cre^/D^b^ animals, we found that infiltration of total CD8 T cells to the brain was unaltered, while virus antigen specific CD8 T cell brain infiltration was reduced (Fig. 5b-d). Inflammatory monocyte infiltration was also reduced in CX3CR1^cre^/D^b^ chimeras (Fig. 6e). These findings suggest that CD8 T cell and inflammatory monocyte responses in the brain are specifically impacted by CNS-myeloid cell antigen presentation.

**Figure 6.**
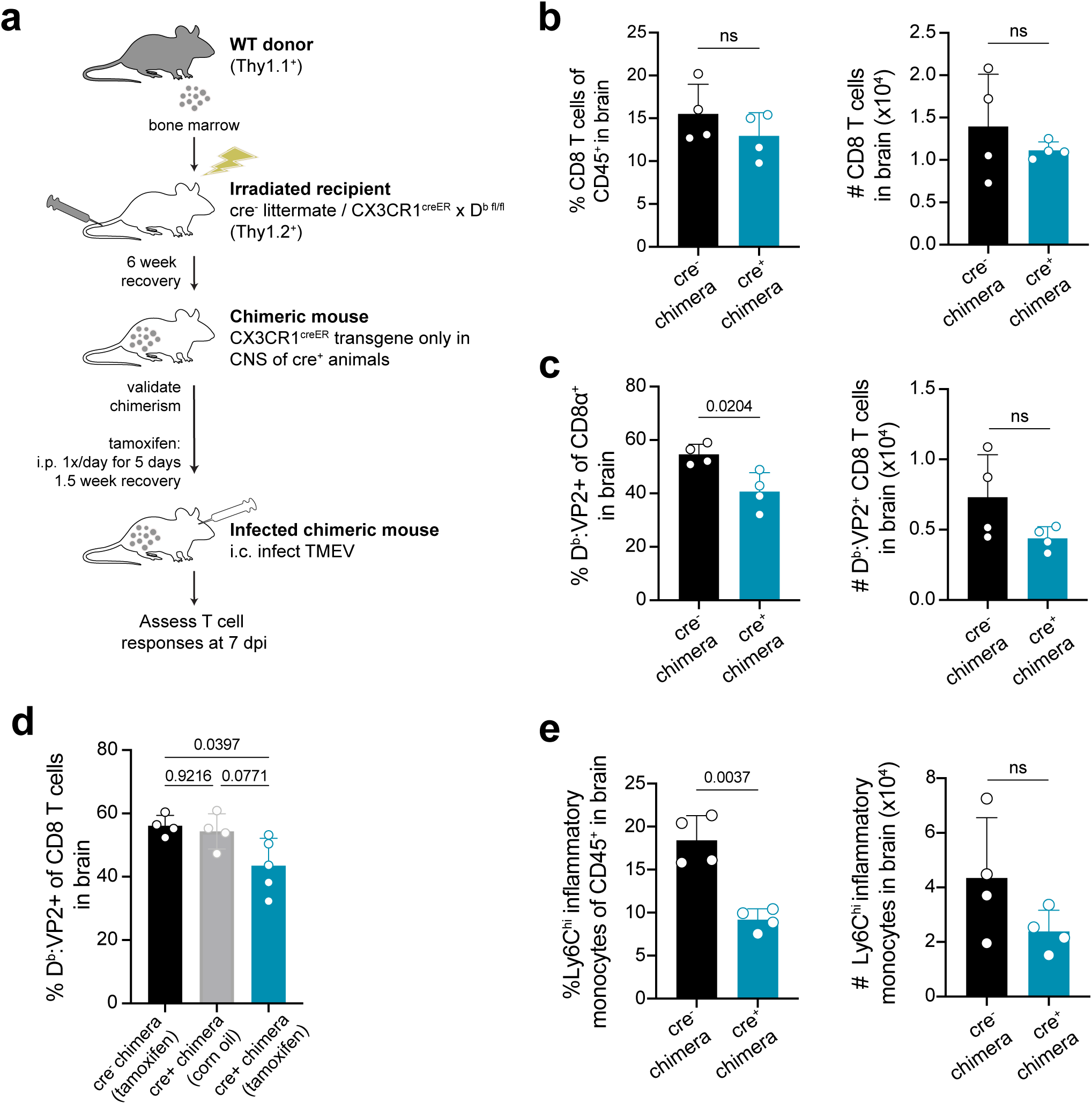
Deletion of H-2D^b^ in only CNS-myeloid cells reduces immune infiltration following TMEV infection. (a) Bone marrow chimeras were generated by lethal irradiation of recipient cre- littermates and CX3CR1^cre^/D^b^ mice followed by reconstitution with WT congenic (Thy1.1) bone marrow. Six weeks later, after validation of chimerism, chimeric mice received intraperitoneal tamoxifen treatment. After recovery, chimeric mice were intracranially infected with TMEV to assess antiviral responses. (b-c) The frequency and number of CD8 T cells (b) and VP2-specific CD8 T cells (c) recovered from the brains of cre^-^ and CX3CR1^cre^/D^b^ mice. (d) A new set of chimeras were generated with the addition of a CX3CR1^cre^/D^b^ group treated with vehicle control. The frequency of D^b^:VP2^+^ CD8 T cells recovered from the brains of these chimeras was quantified. (e) Quantification of Ly6C^hi^ infiltrating inflammatory monocytes recovered from the brains of chimeric cre^-^ and CX3CR1^cre^/D^b^ mice. Data are shown as individual mice with mean from one independent experiment (n=3-6) of ≥ 3 experimental replicates. Error bars represent standard deviation. Two tailed Welch’s *t* test or one-way ANOVA with Tukey’s correction were used to assess statistical significance with ns p ≥ 0.05.

### CNS-myeloid cell antigen presentation to virus-antigen specific CD8 T cells via H-2D^b^ impacts brain atrophy

Given that CNS-myeloid cell deletion of H-2D^b^ resulted in reduced immune infiltration at 7 days post TMEV infection, we next investigated outcomes of viral infection in CX3CR1^cre^/D^b^ animals. We found that CX3CR1^cre^/D^b^ animals experienced reduced weight loss during TMEV infection (Fig. 7a). Next, we measured viral load over the course of infection. TMEV infection is generally cleared between 14-28 days post inoculation^33^, and we determined that CX3CR1^cre^/D^b^ animals were able to clear infectious virus particle with the same kinetics as cre^-^ controls despite reduced immune infiltration (Fig. 7b), indicating that reduced immune infiltration when CNS-myeloid cells lacked expression of H-2D^b^ did not impact clearance of infectious virus.

**Figure 7.**
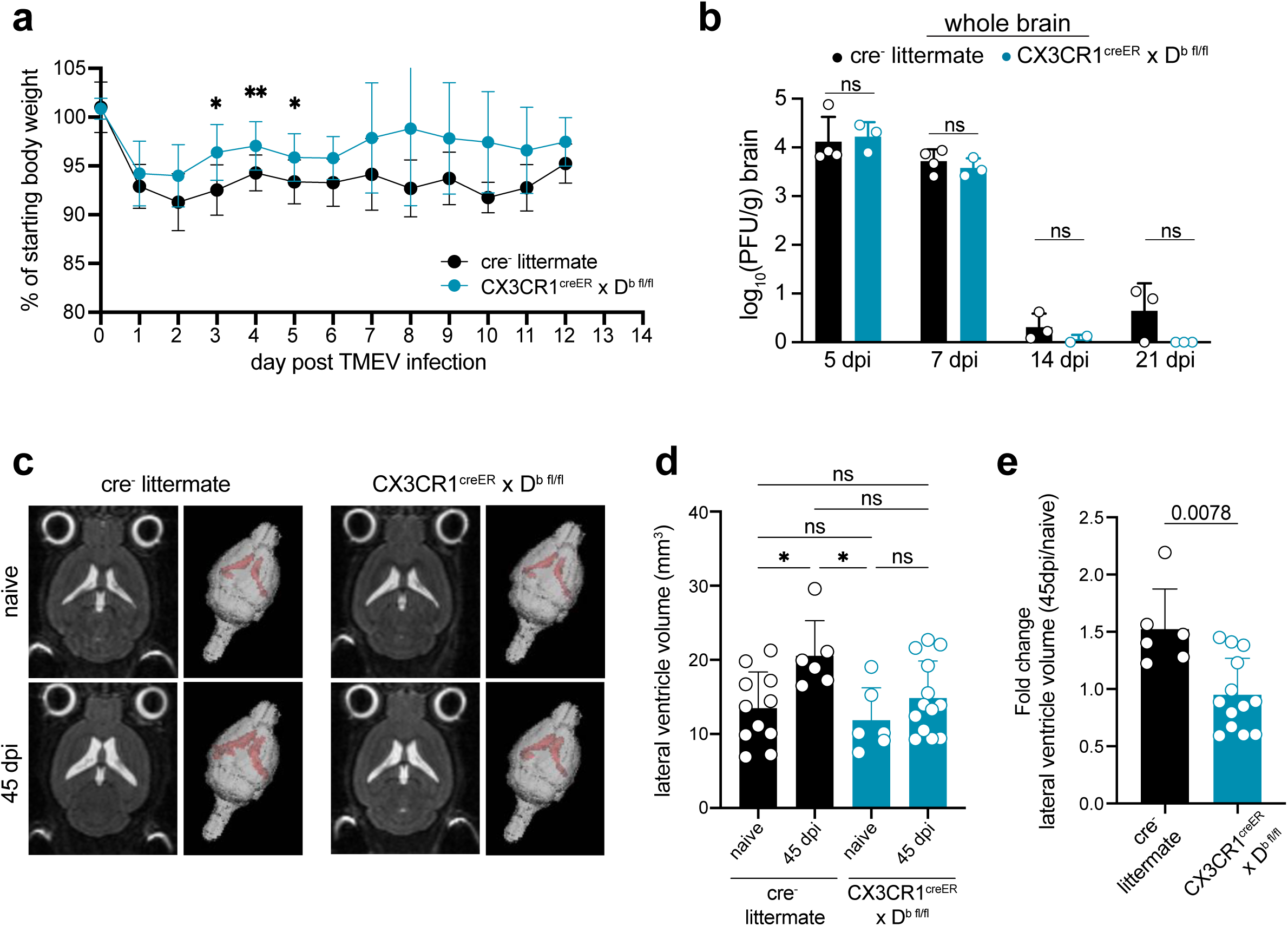
H-2D^b^ expression by CNS-myeloid cells impacts antiviral immunity and brain atrophy. Experimental strategy is identical to Figure 3e, in which tamoxifen administration in CX3CR1^cre^/D^b^ mice and cre^-^ controls is followed by a 6-week reconstitution period before i.c. infection with TMEV. (a) Weight loss over the course of infection is measured in cre^-^ and CX3CR1^cre^/D^b^ mice. (b) Infectious viral load in whole brain homogenates at 5-, 7-, 14-, and 21- days post infection is measured by plaque assay in cre^-^ and CX3CR1^cre^/D^b^ mice. (c) Representative 2D images and 3D renderings of lateral ventricle volumes obtained by T2 weighted MRI of cre^-^ and CX3CR1^cre^/D^b^ mice during naïve conditions and after recovery from TMEV infection (45 dpi). Images were analyzed using Analyze 12.0 (d) Lateral ventricle volume is quantified for naïve and TMEV-recovered cre^-^ controls and CX3CR1^cre^/D^b^ mice. (e) Fold change in lateral ventricle volume from naïve to 45dpi is quantified for TMEV-recovered cre^-^ controls and CX3CR1^cre^/D^b^ mice. Data are shown as individual mice with mean from one independent experiment (n=3-13) of ≥ 2 experimental replicates. Error bars represent standard deviation. Two tailed Welch’s *t* test or one-way ANOVA with Tukey’s correction were used to assess statistical significance with ns p ≥ 0.05.

We next evaluated whether CX3CR1^cre^/D^b^ mice experienced altered neuropathological outcomes resulting from immune responses against TMEV. Cells of the innate immune system infiltrate the brain within 24 hours post TMEV infection and are linked to virus-independent apoptosis of hippocampal neurons and cognitive impairment^50^. Both CX3CR1^cre^/D^b^ and cre^-^ animals that recovered from TMEV infection experienced loss of hippocampal neurons and cognitive impairment measured using the Novel Object Recognition test (NOR) (Sup. Fig. 8), indicating that this outcome was not impacted by CNS-myeloid cell expression of D^b^.

Notwithstanding, reduced immune infiltration could also impact brain atrophy associated with TMEV infection. We have previously shown that brain atrophy, which includes cellular loss, degradation of extracellular matrix, and/or loss of extracellular proteins, is dependent on the D^b^ MHC class I molecule and T cells, suggesting a direct link between CD8 T cell responses and atrophy^51^. Post-infectious brain atrophy was measured by lateral ventricle volume in a T2 weighted MRI. Consistent with previous reports, we found that cre^-^ controls experienced significant brain atrophy at 45 dpi in comparison to naïve cre^-^ counterparts (Fig. 7c-e). CX3CR1^cre^/D^b^ animals did not experience brain atrophy post infection when compared to naïve CX3CR1^cre^/D^b^ controls (Fig. 7c-e). These results indicate that CNS-myeloid cell antigen presentation via H-2D^b^ contributes to the onset of brain atrophy induced by TMEV infection.

## DISCUSSION

The role of local antigen presentation in promoting T cell infiltration of the CNS during neuroinflammation is not entirely understood. Herein, we present a characterization of APCs in the naïve and virally infected brain. We identify CNS-resident myeloid cells, microglia and perivascular macrophages, as local APCs that become activated and upregulate MHC class I molecules during infection. We analyzed the ability of CNS-myeloid cells to promote CD8 T cell trafficking into the brain parenchyma upon viral challenge with both K^b^ or D^b^ restricted antigens, and determined differential requirements for K^b^ and D^b^ expression during acute viral infection. Further, we determined that reducing the antiviral immune response attenuated the development of brain atrophy while still promoting mechanisms of viral clearance, implying that local antigen presentation augments neuropathology in the CNS. Our work provides evidence that CNS-myeloid cells can act as APCs *in vivo* to mediate CNS inflammation.

Microglia and CNS-macrophages play a key role in antiviral defenses during CNS viral infections and can promote T cell effector functions in the CNS^52^. The field has relied heavily on whole cell depletion experiments to assess the contribution of microglia to antiviral defenses. Caution is necessary when interpreting studies employing CSF1R inhibitors to deplete microglia during viral infection, as off-target effects on peripheral immunity were observed in our study and others^53, 54^. This aside, Waltl et al. revealed microglia are key in controlling TMEV propagation and clearance in C57BL/6 mice, positing this was likely due to microglial:T cell interactions^55^. Our data demonstrate that CNS-myeloid cells are positioned to present antigen to modulate antigen-specific CD8 T cells attempting to infiltrate the brain through the BBB during TMEV infection. Importantly, CNS-myeloid cells are not directly infected by TMEV, but may cross-present exogenous antigens such as viral peptides acquired from adjacent infected neurons to impact CD8 T cell infiltration of the CNS during infection^24, 56^. Our work investigating antigen presentation by CNS-myeloid cells during TMEV infection using transgenic mice adds to this body of literature, and extends the possibility that CNS-myeloid cells cross-present antigen derived from infected neurons to modulate CD8 T cell infiltration in response to infection.

MHC class I molecules play multiple roles in immunity, including regulation of CD8 T cell development and cytotoxic activity. In the CNS, MHC class I plays a key role in neuronal development, synaptic plasticity, and axonal regeneration^57^. Nonetheless, in our analysis, CNS-myeloid cells exhibited no obvious development or activation defects upon loss of MHC class I expression. Whether deletion of MHC class I by CNS-myeloid cells during embryogenesis impacts CNS development remains to be studied. MHC class I molecules are also major inhibitory receptors for NK cells, and lack of expression can trigger NK-mediated killing^58^. However, our data indicate that CNS-myeloid cells lacking K^b^ and D^b^ were not reduced in population and hence are not killed by NK cells. This is in contrast to what has been reported for *β*2m floxed mice, a model that ablates H-2K^b^/H-2D^b^ as well as nonclassical MHC class I molecules and can impact NK cell licensing and recognition^56^. We therefore contend that our cre-lox strategy specific for K^b^ and D^b^ provides a means to study discrete contributions of each MHC class I molecule in isolation without off-target effects from Class Ib molecules.

The data presented herein demonstrate that CNS-myeloid cells are required for immune infiltration of the brain in a context-dependent manner. The difference between K^b^ and D^b^ may be explained by the importance of the H-2D^b^ molecule in driving CD8 T cell recognition of the immunodominant epitope of TMEV. However, the engineered TMEV-OVA virus experiences an attenuation in virulence as compared to the wt virus^34^, potentially impacting our results. Nonetheless, we posit that K^b^/D^b^ differences extend beyond CD8 T cell recognition of TMEV/TMEV-OVA and infer differences between K^b^ and D^b^ during neuroinflammation. We found that K^b^ and D^b^ were expressed in differing levels by immune cells in the naïve and infected CNS. The H-2K and H-2D genes are differentially expressed by unique subsets of neurons in the developing and mature CNS, indicating different functionality^59^. Whether K^b^ and D^b^ differentially impact antigen presentation during other neuroinflammatory conditions remains to be studied.

In this study, CNS-myeloid cell antigen presentation impacted antiviral CD8 T cell and inflammatory monocyte infiltration of the brain. CD8 T cells have the ability to secrete chemokines that attract pathogenic monocytes during lymphocytic choriomeningitis virus (LCMV) infection^60^. Our findings contend that recruitment of monocytes to the brain is augmented by antigen presentation and CD8 T cell responses, and CD8 T cells may produce chemoattractants such as CCL2 to promote monocytic brain infiltration^61^. Despite this, CNS-myeloid cell antigen presentation did not alter hippocampal neuron damage, perhaps due to intact early innate immune processes with the capacity to contribute to neuronal loss^50^. Mice with defects in CNS-myeloid cell antigen presentation were however protected from brain atrophy. Our working model is that damage to hippocampal neurons occurs within the first few days as a result of innate immune processes, while local antigen presentation via H-2D^b^ drives CD8 T cells that contribute to brain atrophy in regions outside of the hippocampus. It is therefore conceivable that brain atrophy can be attenuated by reducing antiviral CD8 T cell infiltration, and CD8 T cell infiltration is likely mediated by local antigen presentation by CNS-myeloid cells. Given the recent link between CD8 T cells and neurodegenerative conditions such as Alzheimer’s disease and Parkinson’s disease, our findings are highly relevant to neurologic disease^8,9,10^.

Our findings suggest that additional APCs must play a role in antiviral immune infiltration of the CNS. Meningeal APCs can promote CD4 T cell infiltration of the inflamed CNS^62^. However, meningeal antigen presentation to CD8 T cells remains to be investigated. Critical APCs may also include glial cells, such as oligodendrocyte precursor cells (OPCs), which cross present antigen via MHC class I when exposed to IFN*γ*during demyelination^63^. The contribution of additional APCs to promoting CD8 T cell infiltration of the inflamed CNS will be the topic of further investigation.

The classical view of antiviral T cell responses posits that once activated CD8 T cells search for cognate peptide presented on MHC class I by infected cells after infiltrating the tissue. While once controversial, direct CD8 T cell engagement with neurons during neuroinflammation is becoming more widely accepted and may play a critical role in pathogen clearance. CD8 T cells forming immune synapses with TMEV infected neurons^64^. Similarly, latent herpes simplex virus 1 (HSV) reactivation is modulated by CD8 T cells recognizing cognate antigen presented by neuronal MHC class I^5^. Finally, neuronal antigen presentation is required for CD8 T cell mediated clearance of the *Toxoplasma gondii* parasite in multiple phases of infection^65^. It is unknown whether neuronal MHC class I plays a key role in CD8 T cell containment of TMEV or TMEV-OVA, though these studies are ongoing. Overall, our results suggest that CNS-myeloid cells play a role in promoting CD8 T cell infiltration of the infected CNS in an antigen-dependent manner, but compensatory mechanisms exist that promote viral clearance. Thus, MHC class I expression on multiple CNS cell types may be required for unique immunological mechanisms that promote CD8 T cell infiltration through brain barriers and CD8 T cell mediated viral clearance in a cell-specific manner.

In conclusion, our study sheds light on the importance of CNS-myeloid cell antigen presentation during CNS viral infection and reveals marked differences between the H-2K^b^ and H-2D^b^ MHC class I molecules. These findings have important implications for our understanding of CD8 T cell mediated antiviral immunity and immunopathology in the CNS, as considerable focus has been exclusively placed on antigen presentation by the H-2K^b^ molecule due to the wide availability of reagents and model antigens. Our study also emphasizes the importance of MHC class I antigen presentation in the CNS in the context of neurodegeneration. Further investigation of the roles of specific MHC class I alleles and antigen presentation by discrete cell types will be crucial, especially given increasing literature linking CD8 T cell responses to aging and neurodegenerative diseases. These studies should offer new insights into the therapeutic potential of targeting CNS infiltration of antigen-specific T cells as a means to attenuate T cell mediated brain atrophy in neurological disease^8,9,10, 66^.

## ACKNOWLEDGEMENTS

We would like to thank Drs. Charles Howe and LongJun-Wu for their crucial help in preparing this manuscript and sharing resources. We would like to thank Manling Xie, Jiaying (Fiona) Zheng, and Dr. Upkong Eyo for their technical assistance and help in designing experiments. We would also like the thank the NIH tetramer core facility for providing tetramers that were used in this work.

## AUTHOR CONTRIBUTIONS

ENG, CEF, CGL, KA, designed and performed the majority of experiments. LTY, CSM, RHK, ZPT, FJ, MJH performed some experiments and/or provided feedback towards experimental design and manuscript conceptualization. ENG and AJJ wrote the manuscript. All authors have read the manuscript and provided essential feedback, and all co-authors agree with the data presented.

---

**Supplemental Figure 1.**
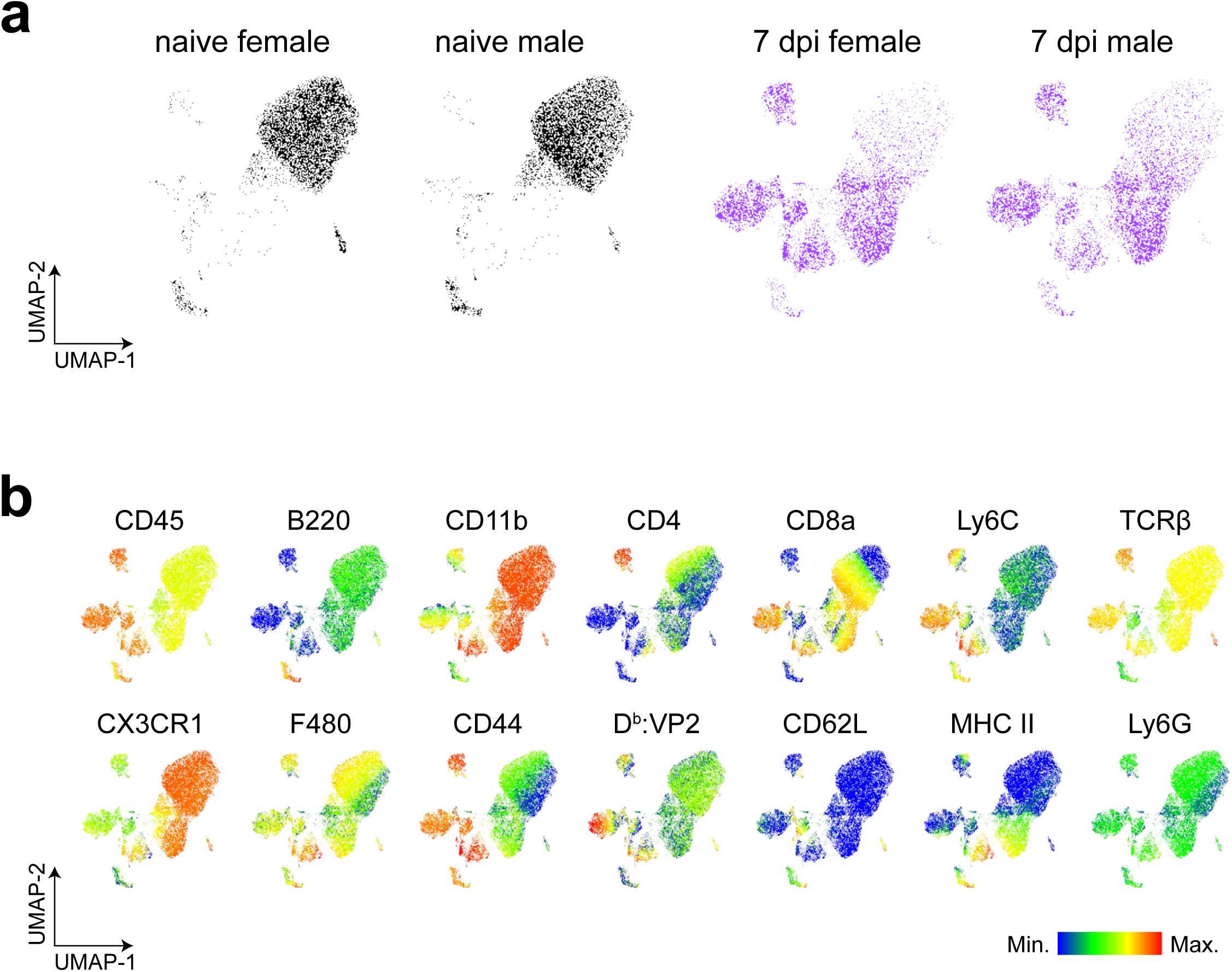
Analysis of immune compartment in naïve and TMEV-infected mice. Related to Figure 1. (a) UMAP visualization of the CD45^+^ cell subsets isolated from the brain in Figure 1, color-coded by infection status and biological sex. (b) UMAP visualization of the CD45^+^ cell subsets isolated from the brain, colored by expression of phenotypic cell surface markers used in flow cytometry analysis. Data are representative of ≥ 3 independent experiments.

**Supplemental Figure 2.**
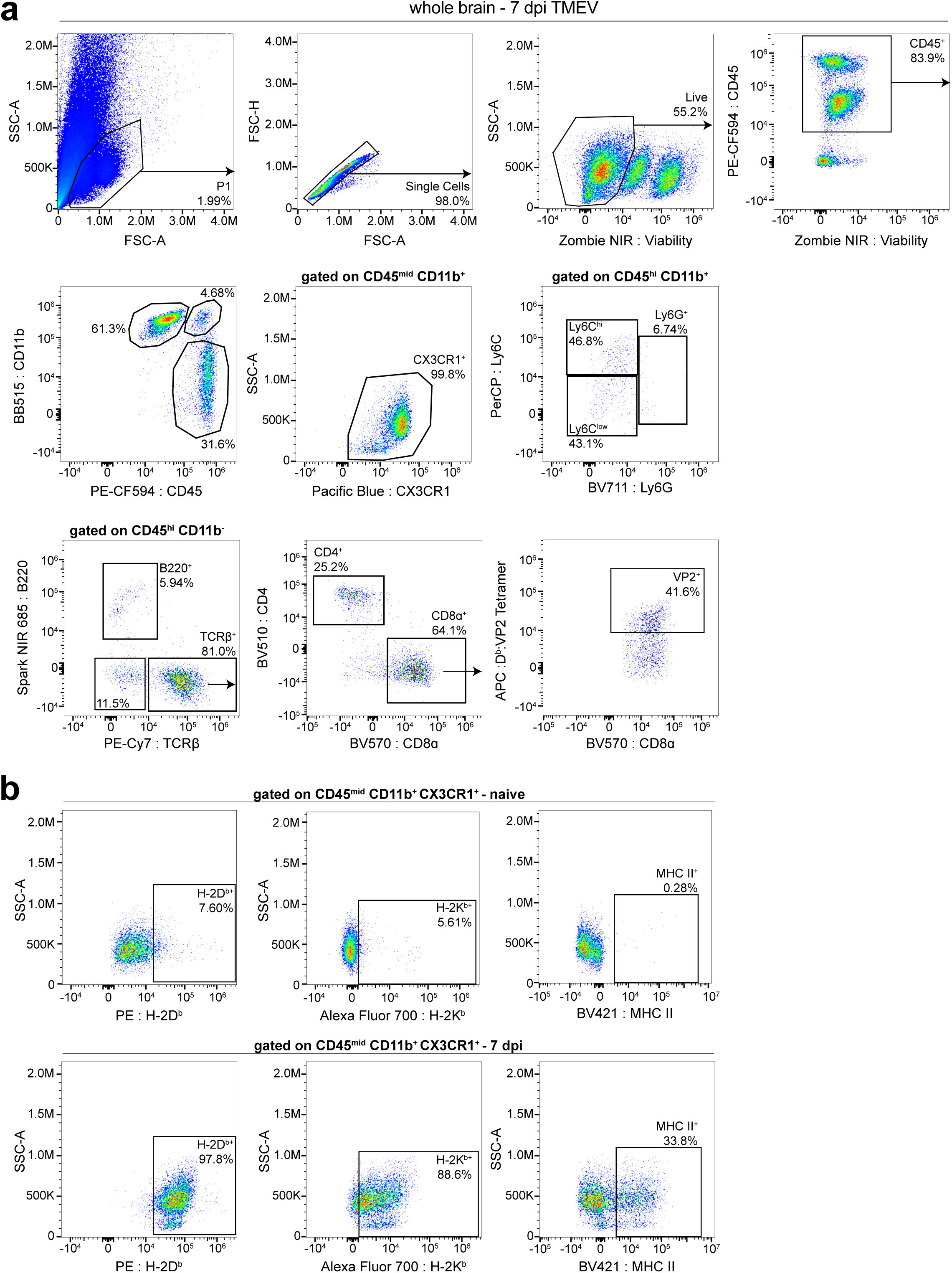
Gating strategies for flow cytometry. (a) Representative flow cytometry gating strategies for manual analysis of immune cells in whole brain samples. Plots are representative of a TMEV infected mouse for ease of viewing. (b) Representative gating for analysis of MHC class I and MHC class II expression on CNS-myeloid cells. Gates were drawn using Fluorescence Minus One (FMO) controls.

**Supplemental Figure 3.**
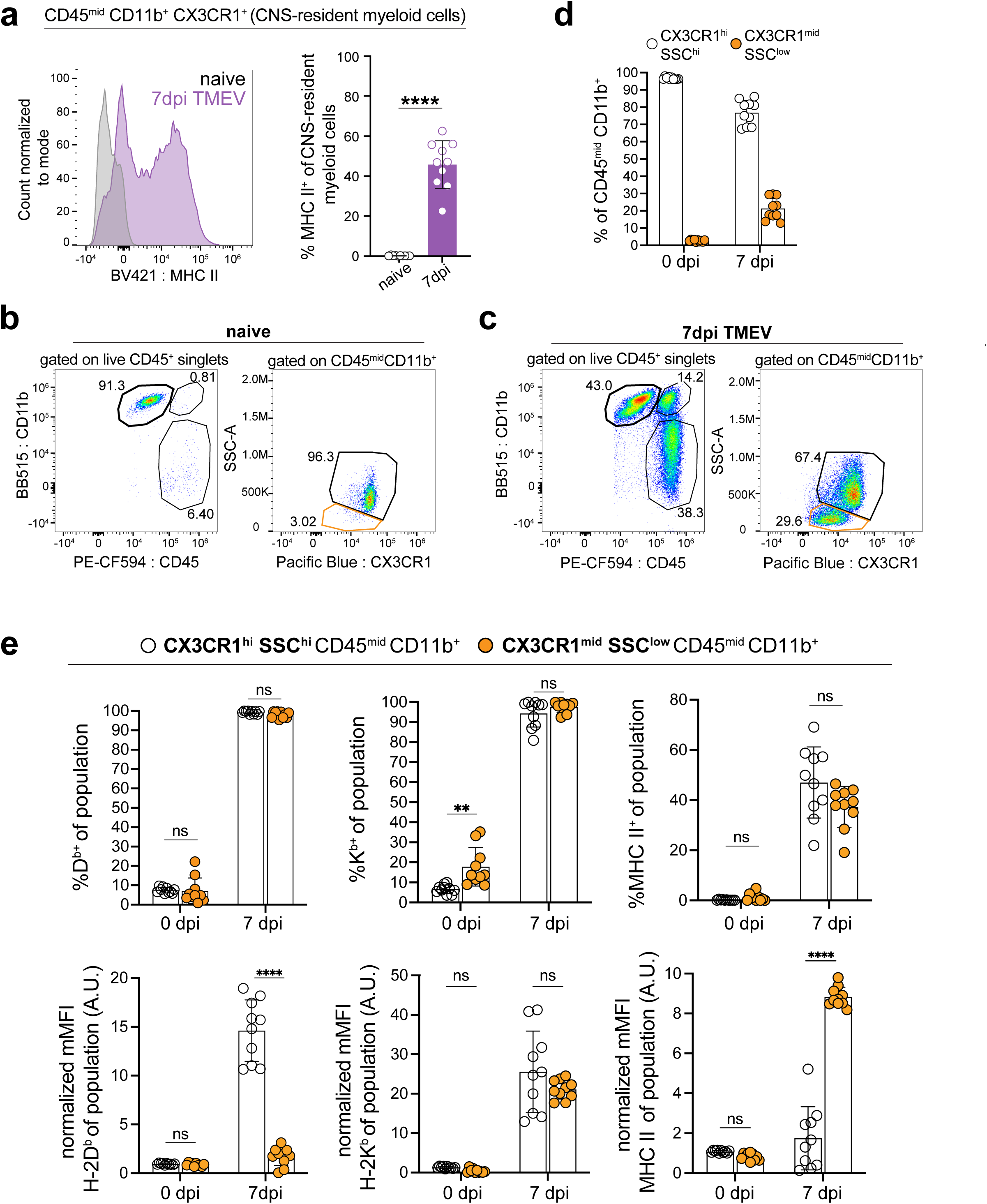
TMEV infection results in activation of CNS-resident myeloid cells and expression of MHC class I/II. Related to Figure 2. (a) MHC class II expression on CNS- myeloid cells during naïve and steady state conditions, demonstrated by representative histogram. The frequency of MHC II positive cells and normalized MFI of MHC II on the population is quantified (b) Representative flow cytometry plots of CNS-myeloid cells (CD45^mid^ CD11b^+^ CX3CR1^+^) in the naïve state. (c) Representative flow cytometry plots of CNS-myeloid cells (CD45^mid^ CD11b^+^ CX3CR1^+^) at 7 days post TMEV infection. (d) Quantification of CX3CR1^hi^ SSC^hi^ and CX3CR1^mid^ SSC^mid^ CNS-myeloid cells during naïve and TMEV infected conditions. (e) Expression of H-2Db, H-2Kb, and MHC II on CX3CR1^hi^ SSC^hi^ (granular) and CX3CR1^mid^ SSC^mid^ is quantified by percent-positive of population and normalized median fluorescence intensity. Data are representative of ≥ 3 independent experiments and presented as mean +/- SD, one-way ANOVA with Tukey’s correction was used to assess statistical significance with ns p ≥ 0.05. dpi, days post injury.

**Supplemental Figure 4.**
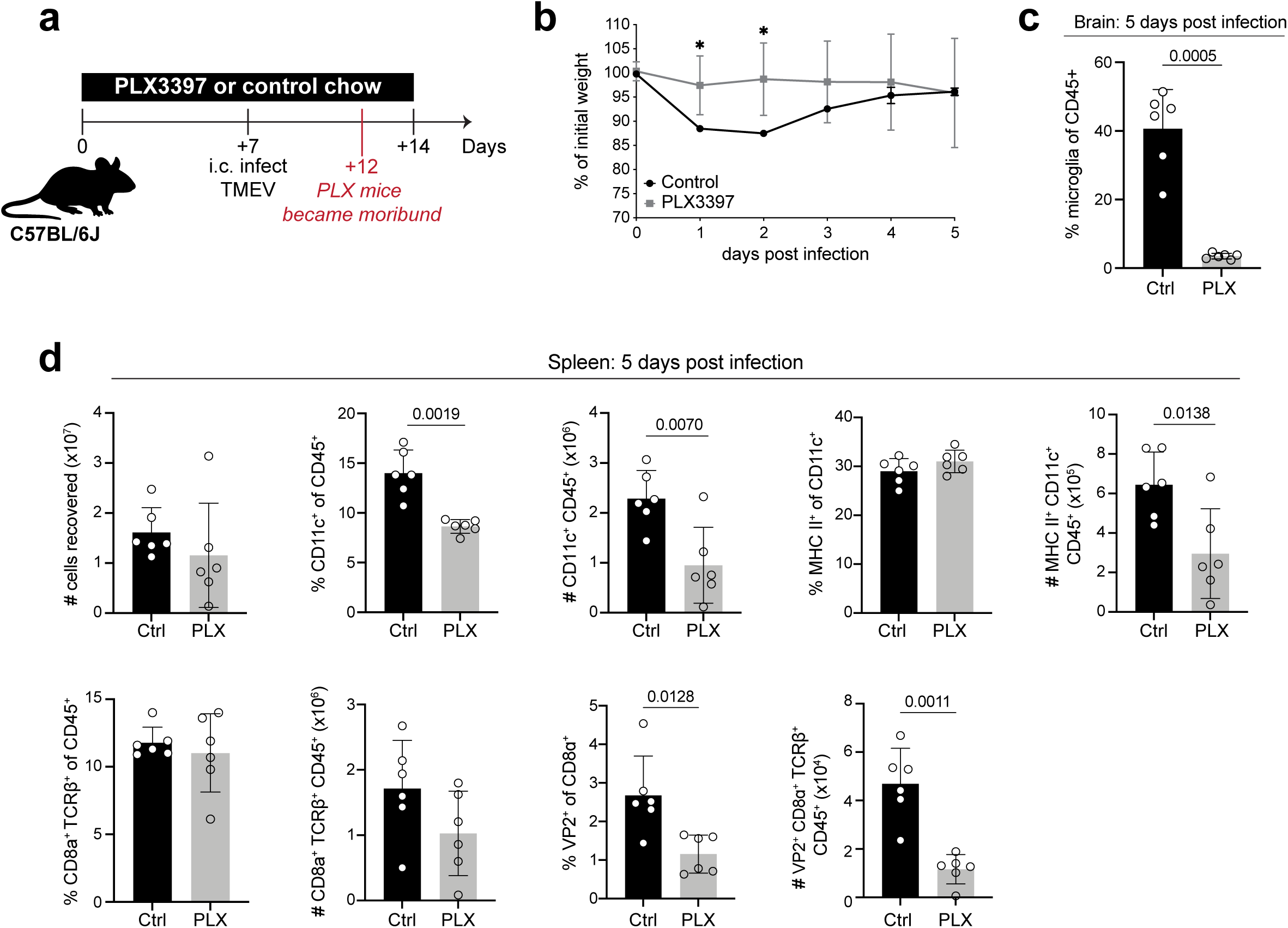
Use of a CSF1R inhibitor reduces peripheral myeloid populations responsible for T cell priming. (a) Experimental timeline. C57BL/6 female mice were fed chow containing PLX3397 or control chow for 7 days prior to infection with TMEV and were continuously fed PLX throughout the course of infection. Flow cytometry analysis was executed when infected PLX treated mice became moribund at 5 dpi TMEV. (b) Weight loss as a result of infection was measured in control and PLX3397 treated mice. (c) The frequency of CNS-myeloid cells in the brain was quantified using flow cytometry in control and PLX3397 treated animals. (d) Analysis of splenic immune populations at 5 dpi, including: total cell number, frequency and number of CD11c^+^ cells, frequency and number of MHC II^+^ CD11c^+^ cells, frequency and number of CD8 T cells, and frequency and number of virus specific (D^b^:VP2^+^) CD8 T cells. Data are presented as mean +/- SD, significance was assessed using 2-tailed unpaired Student’s *t* test with ns p ≥ 0.05.

**Supplemental Figure 5.**
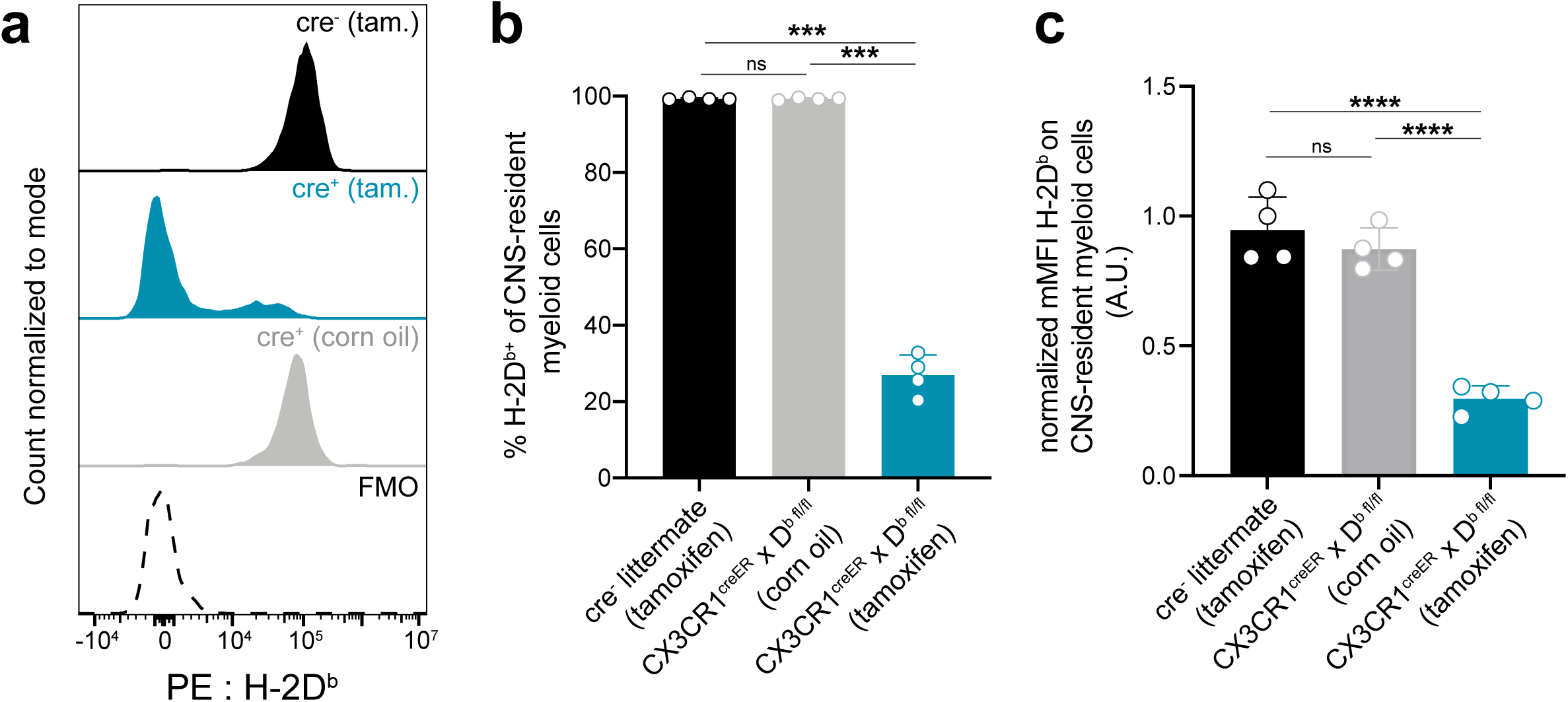
No spontaneous recombination of floxed H-2D^b^ transgene occurs in CX3CR1^cre^/D^b^ mice. Related to Figure 3. CX3CR1^cre^/D^b^ animals were treated with vehicle control (corn oil) and compared to tamoxifen-treated cre^+^ and cre^-^ animals. (a) Representative histogram of H-2D^b^ expression on CNS-myeloid cells. (b) Quantification of percent D^b^ positive CNS-myeloid cells at 7 dpi TMEV (c) Normalized MFI of H-2D^b^ on CNS-myeloid cells. Data are shown as individual mice with mean from one independent experiment (n=4) of ≥ 2 experimental replicates. Error bars represent standard deviation. One-way ANOVA with Tukey’s correction was used to assess statistical significance with ns p ≥ 0.05.

**Supplemental Figure 6.**
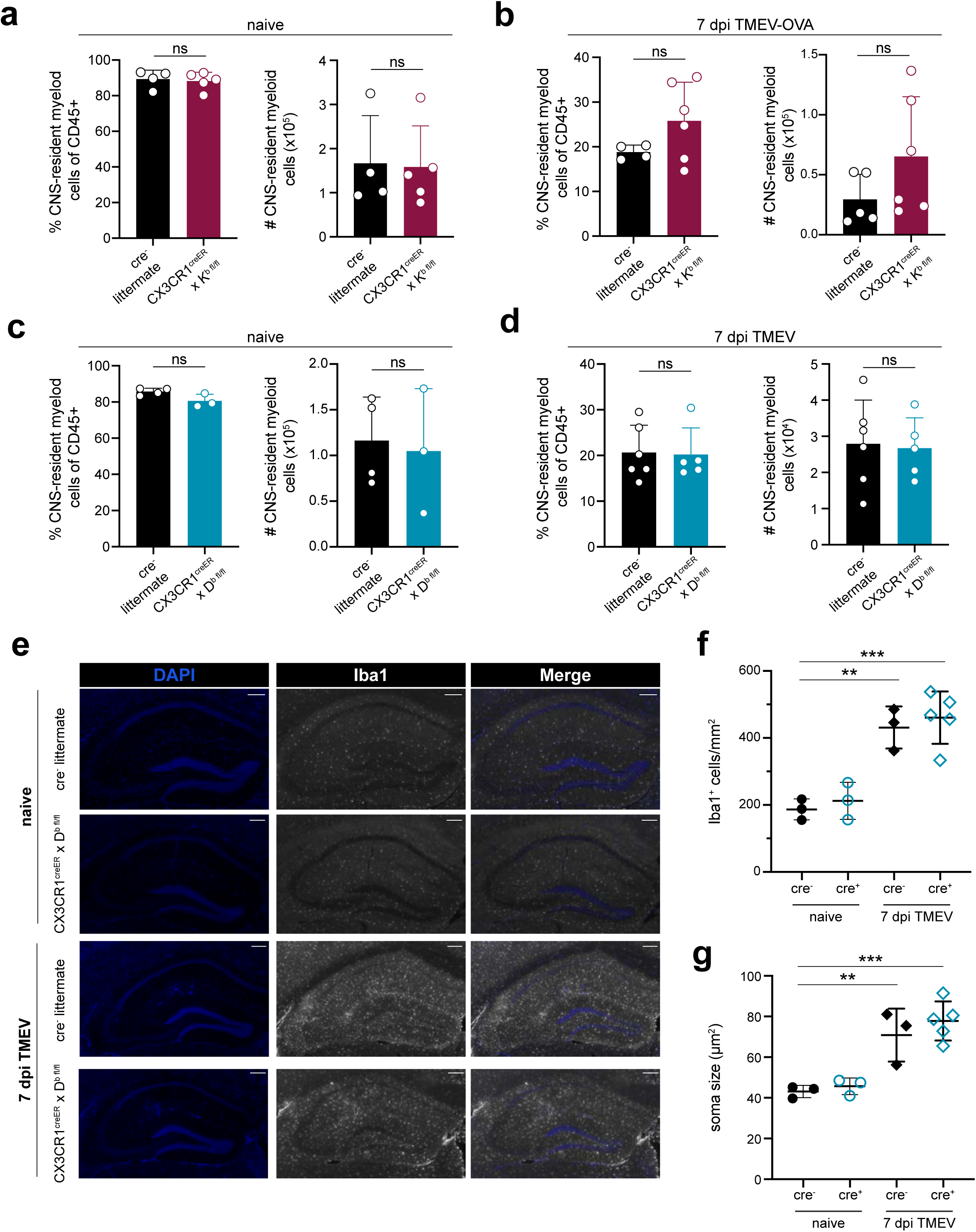
No cell-intrinsic defects detected when CNS-myeloid cells lack MHC class I. Related to Figure 3. (a-b) Flow cytometic quantification of CNS-myeloid cells isolated from cre^-^ and CX3CR1^cre^/K^b^ mice during naïve conditions (a) and at 7 dpi with TMEV-OVA (b). (c-d) Flow cytometic quantification of CNS-myeloid cells isolated from cre^-^ and CX3CR1^cre^/D^b^ mice during naïve conditions (c) and at 7 dpi with TMEV (d). (e) Hippocampal sections collected at 0 and 7 days post TMEV infection were immunostained for Iba1 (white). Representative images are shown. (f) The density of hippocampal Iba1+ cells/mm^2^ is quantified for the groups shown in (e). (g) The soma size of hippocampal Iba1+ cells were measured for the groups shown in (e). For flow cytometry experiments, CNS-myeloid cells are defined as live, single cells, expressing CD45^mid^CD11b^+^CX3CR1^+^. Flow data are shown as individual mice with mean from one independent experiment (n=3-6) of at least three experimental replicates. Immunofluorescence data are shown as representative individual mice with mean from one independent experiment (n=3-5) of at least two experimental replicates. Iba1+ cell density is quantified from two hippocampal sections and plotted as average per mouse. Iba1+ cell soma size is quantified from 25 microglia per mouse and plotted as average per mouse. Error bars represent standard deviation. Two tailed Welch’s *t* test or one-way ANOVA with Tukey’s correction were used to assess statistical significance with ns p ≥ 0.05.

**Supplemental Figure 7.**
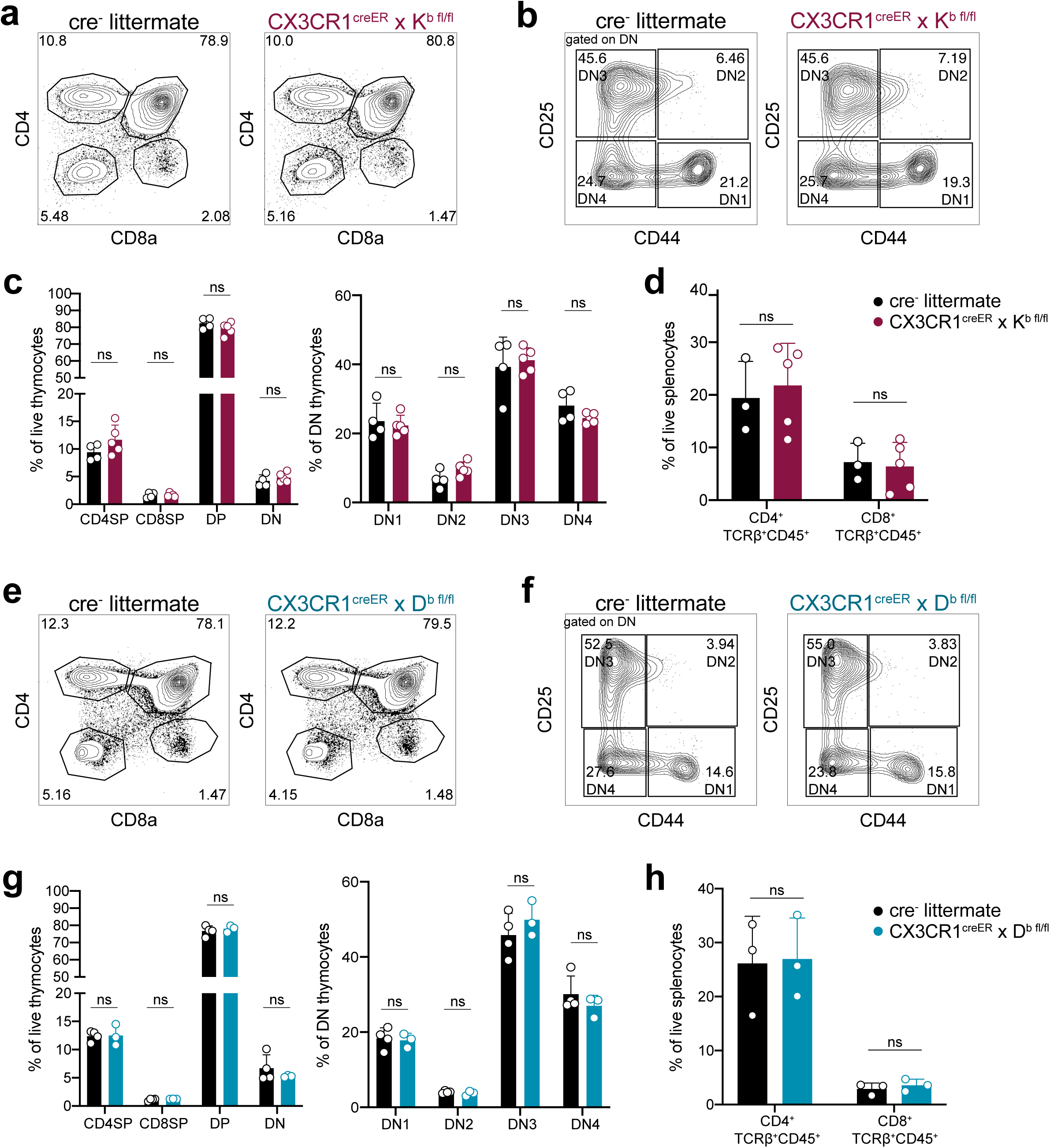
Mice lacking MHC class I on CNS-myeloid cells have normal T cell development after tamoxifen treatment. Representative thymic gating strategies for tamoxifen treated cre^-^ littermates and CX3CR1^cre^/K^b^ are shown in (a) and (b). After gating on single, live, CD45+ cells, CD4 and CD8 were used to measure double negative (DN), double positive (DP), CD4 single positive (CD4SP), and CD8 single positive (CD8SP) (a). (b) Within the DN gate, DN1- 4 were defined and quantified as follows: CD44+CD25- (DN1), CD44+CD25+ (DN2), CD44- CD25+ (DN3), and CD44-CD25- (DN4). (c) Frequencies of CD4SP, CD8SP, DP, DN, and DN1- 4 are quantified between cre^-^ and CX3CR1^cre^/K^b^ animals. (d) CD4 and CD8 T cell frequencies were quantified in the spleens of cre^-^ and CX3CR1^cre^/K^b^ mice. (e-h) An identical analysis of T cell development was performed on cre^-^ littermates and CX3CR1^cre^/D^b^ animals. (e-f) Representative images and quantification of T cell subsets are shown. Data are represented as individual mice with mean from one independent experiment (n=3-5) out of at least three experimental replicates. Error bars represent standard deviation. Two tailed Welch’s *t* test was used to assess statistical significance, with ns p ≥ 0.05.

**Supplemental Figure 8.**
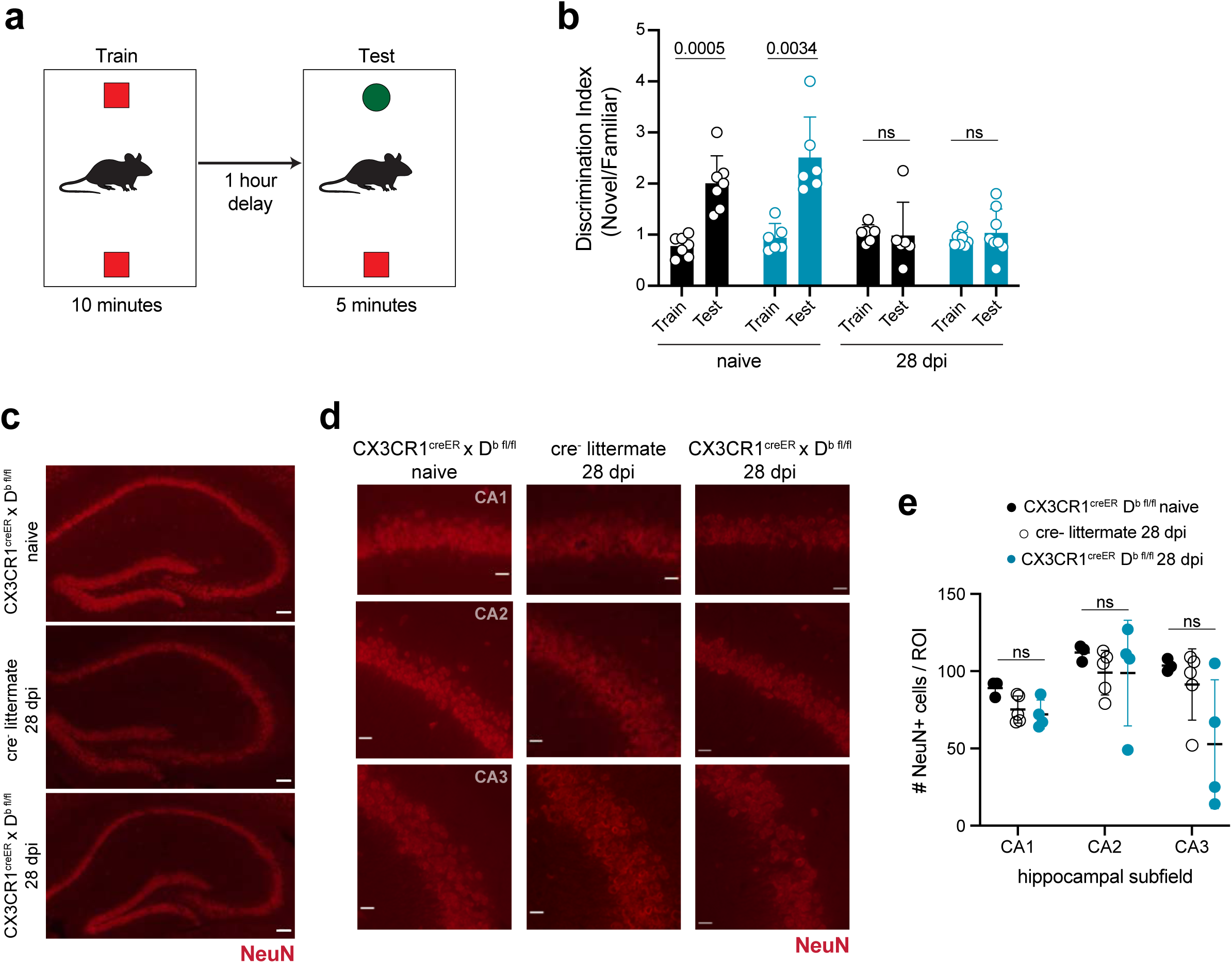
Deletion of D^b^ from CNS-myeloid cells does not prevent cognitive deficits and damage to hippocampal neurons resulting from TMEV infection. Related to Figure 7. (a) Experimental paradigm for the Novel Object Recognition test of learning and recognition memory. After habituation, mice are exposed to two objects during the training period. After a delay, a novel object replaces a familiar object. Investigation of each object is quantified during both training and testing phases. (b) Discrimination index, or number of sniffs of novel object as compared to familiar object, in cre^-^ and CX3CR1^cre^/D^b^ mice under naïve conditions or after recovery from TMEV. (c-d) NeuN staining of the hippocampus in naïve and TMEV recovered control and CX3CR1^cre^/D^b^ animals. Equivalent ROIs in CA1, CA2, and CA3 were drawn and the number of NeuN+ cells were counted in each ROI. (g) Quantification of NeuN+ cells per ROI in each hippocampal area in recovered cre^-^ and CX3CR1^cre^/D^b^ mice compared to naive CX3CR1^cre^/D^b^ mice. Data are represented as individual mice with mean from one independent experiment (n=3-8) out of at least two experimental replicates. Error bars represent standard deviation. Two tailed Welch’s *t* test or one-way ANOVA with Tukey’s correction were used to assess statistical significance, with ns p ≥ 0.05.

